# JAK/STAT pathway promotes *Drosophila* neuroblast proliferation via the direct *CycE* regulation

**DOI:** 10.1101/2020.07.10.195875

**Authors:** Lijuan Du, Jian Wang

## Abstract

How neural stem cells regulate their proliferative potential and lineage diversity is a central problem in developmental neurobiology. *Drosophila* Mushroom bodies (MBs), centers of olfactory learning and memory, are generated by a specific set of neuroblasts (Nbs) that are born in the embryonic stage and continuously proliferate till the end of the pupal stage. Although MB presents an excellent model for studying neural stem cell proliferation, the genetic and molecular mechanisms that control the unique proliferative characteristics of the MB Nbs are largely unknown. Further, the signaling cues controlling cell cycle regulators to promote cell cycle progression in MB Nbs remain poorly understood. Here, we report that JAK/STAT signaling pathway is required for the proliferation activity and maintenance of MB Nbs. Loss of JAK/STAT activity severely reduces the later-born MB neuron types and leads to premature neuroblast termination, which can be rescued by tissue-specific overexpression of *CycE* and *diap1*. Higher JAK/STAT pathway activity in MB results in more neurons, without producing supernumerary Nbs. Furthermore, we show that JAK/STAT signaling effector Stat92E directly regulates *CycE* transcription in MB Nbs. Finally, MB Nb clones of loss or excess *CycE* phenocopy those of decreased or increased JAK/STAT signaling pathway activities. We conclude that JAK/STAT signaling controls MB Nb proliferative activity through directly regulating *CycE* expression to control cell cycle progression.

## Introduction

During brain development, neural stem cells (NSC) produce lineages of post-mitotic neurons or glia through spatially and temporally controlled self-renewing cell divisions. Like other stem and progenitor cells, an NSC divides asymmetrically to form two distinct daughter cells that differ in fate. One daughter cell retains all features of a stem cell and continues to proliferate, whereas the other daughter cell enters the path of differentiation. Deregulation of NSC proliferation can lead to brain tumor or to a premature depletion of the progenitor pool. Importantly, both the mode of cell division and the controlling mechanisms of progenitor proliferation and differentiation, remain highly conserved between vertebrates and invertebrates (Gage and Temple, 2013) (Sousa-Nunes et al., 2010) (Homem and Juergen A Knoblich, 2012).

The *Drosophila* nervous system has been shown to be an extremely valuable model for studying the regulating mechanisms of stem cell asymmetric division (Jörg Betschinger and Jürgen A Knoblich, 2004) (Yu et al., 2006) (Doe, 2008) (Gómez-López et al., 2014). *Drosophila* neural stem cells, called neuroblasts (Nbs), divide asymmetrically to produce two daughter cells. The larger daughter cell self-renews, whereas smaller daughter cell called the ganglion mother cell (GMC) divides once more and differentiates into neurons or glia. During Nb asymmetric division, a conserved apical protein complex consisting of Par-3/Bazooka, Par-6, and aPKC is selectively partitioned into Nb to maintain the Nb fate, and on the other side, basal proteins such as Prospero, Miranda and Brain Tumor are recruited into the GMC to confer the differentiation fate (C.-Y. Lee et al., 2006a) (Joerg Betschinger et al., 2006) (Bello et al., 2006) (C.-Y. Lee et al., 2006b).

The proliferative potential and the timing for cell cycle exit or termination of Nbs have to be tightly controlled to generate diverse types of neurons and glia. The *Drosophila* larval brain contains ∼200 Nbs. Most of them terminate before late larval stages and generate only a few dozen neurons. However, 4 mushroom body neuroblasts (MB Nbs) exhibit uninterrupted proliferation until the late pupal stages, and each generates hundreds of neurons of three subtypes: γ, α’/β’, and α/β neurons in succession (Ito et al., 1997) (T. Lee et al., 1999). The exceptional proliferation activity of MB Nbs provides a unique model to investigate the molecular and genetic mechanisms controlling the persistent neural progenitor proliferation, which are largely unknown. Further, the regulatory mechanisms linking MB Nb proliferation to cell-cycle regulators remain elusive.

The MARCM-based genetic screen has facilitated us to identify multiple genes required for different aspects of MB development (T. Lee and Luo, 1999) (J. Wang et al., 2002) (Zheng et al., 2003) (J. Wang et al., 2006). Via a similar genetic screen, we discovered that the JAK/STAT signaling pathway is required to promote cell-cycle progression and prevent premature termination of MB Nbs, but it does not affect cell fate differentiation of MB neurons. Furthermore, we demonstrated that JAK/STAT signaling mediates MB Nb cell-cycle progression mainly through the direct activation of *Cyclin E* (*CycE*), an essential G1 cyclin rate-limiting for progression into S phase (Resnitzky et al., 1994). Interestingly, *CycE* is also a primer target of Hippo (Hpo)/Yorkie (Yki) pathway (Wu et al., 2008). We found that JAK/STAT and Yki function in parallel, and both function through *CycE* and *diap1* to sustain Nb proliferation. These results for the first time revealed the essential and parallel roles of two major cell proliferation controlling signaling pathways JAK/STAT and Hpo/Yki pathways in regulating MB neurogenesis, and provided a regulatory link between Nb proliferation and transcriptional regulation of critical G1/S cell cycle regulator, *CycE*.

## Results

### Lack of JAK/STAT signaling activity leads to defects in MB neurogenesis

A MB Nb sequentially generates about 500 neurons of three subtypes: γ, α’/β’, and α/β through undisrupted asymmetrical division from the embryonic stage to the end of pupal stage. In the adult brains, the earliest born γ neurons have one major medially projecting axon branch, while the later born α’/β’ and α/β neurons have two axon branches that project dorsally and medially, respectively (Ito et al., 1997) (T. Lee et al., 1999) (Fig. 1A). The morphologically well distinguishable neuron subtypes make the phenotypic analysis of MB neurogenesis considerably easy (Fig. 1A). The MARCM clones of MB Nb generated at early developmental stages and labeled with *GAL4-OK107*>*UAS-mCD8-GFP* allow us to visualize the gross morphology of the MB and to follow one Nb through different developmental stages (T. Lee and Luo, 1999). When it was created at newly hatched larvae (NHL), a wild type MB Nb generated all three subtypes of neurons (Fig. 1B). However, a MB Nb mutant for JAK/STAT pathway components the receptor *domeless* (*dome*) or the JAK kinase *hopscotch* (*hop*) produced much fewer neurons and most of the neurons were of γ neuron morphology (γ-only phenotype) (Fig. 1C-E,H). The *dome* mutant phenotype was fully rescued by overexpression of *Dome* or *Stat92E*^*ΔNΔ*^*C*, a dominant active form of the JAK/STAT pathway downstream effector Stat92E (Ekas et al., 2010) (Fig. 1F-H), indicating that this phenotype is caused by loss of JAK/STAT signaling activity.

**Figure 1.**
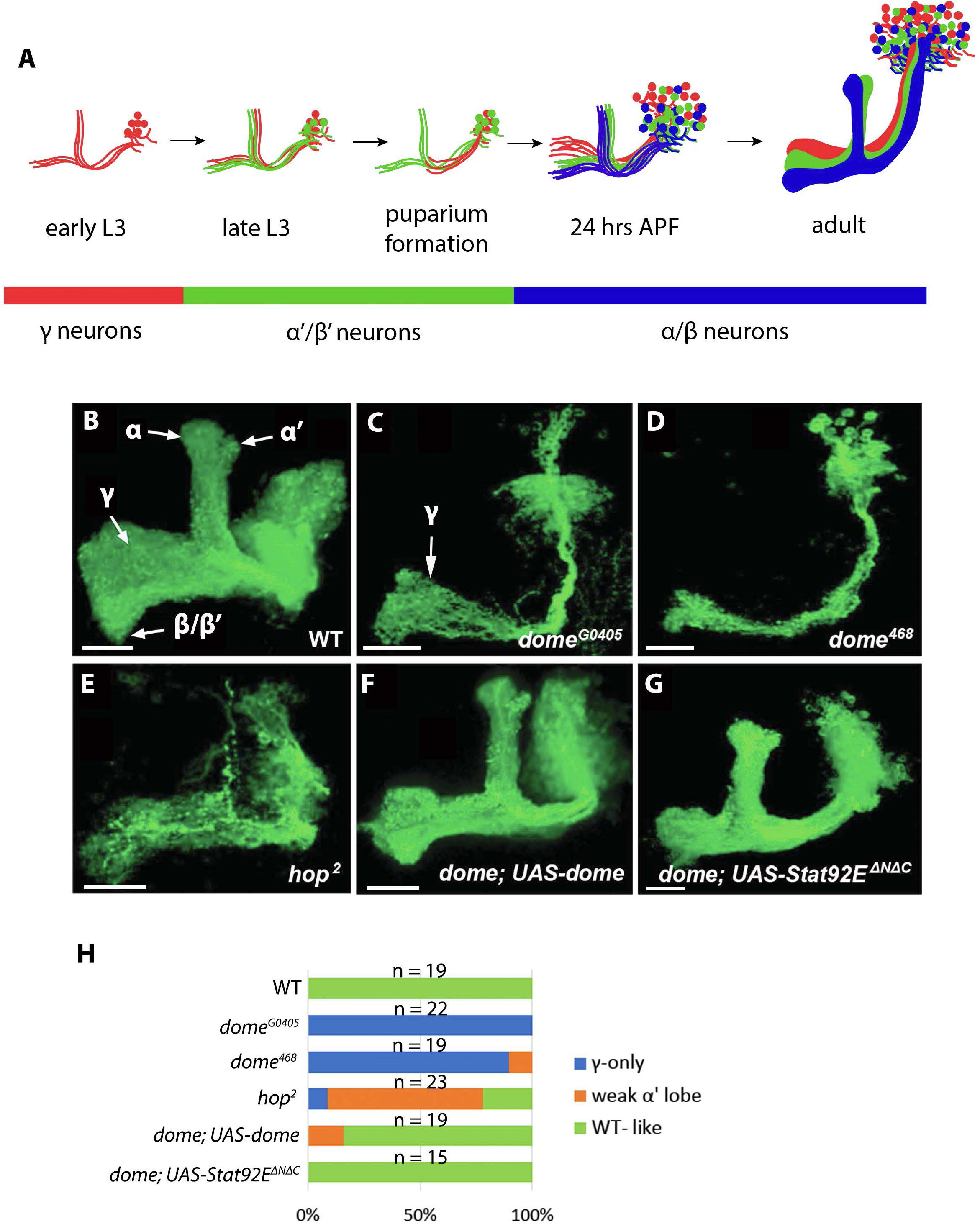
JAK/STAT signaling pathway is required for the MB neurogenesis. **(A)** Schematic summary of MB neurogenesis, depicting the three subtypes of neurons generated from a common Nb precursor in succession: γ (red), α’/β’ (green) and α/β (blue) neurons. Note larval specific γ neurons were pruned and remodeled at the time of puparium formation; different neuron types have distinct axonal projection patterns in the adult brain. APF, after puparium formation. **(B-G)** Representative confocal images of mushroom body neuroblast (MB Nb) clones induced in the newly hatched larvae and examined in the adult brains. All clones were labeled by *GAL4-OK107>mCD8-GFP*. **(B)** WT MB Nb clone, midline of the brain is at the left side of the image in this and all subsequent images. Five axon lobes of three neuron subtypes, γ, α, α’, β, and β’ lobe, were labeled with arrows. **(C)** *dome*^*G0405*^ MB Nb clone, note the reduction in neuron numbers and the γ-only phenotype. **(D)** *dome*^*468*^ MB Nb clone, showing phenotype similar to that of *dome*^*G0405*^. **(E)** *hop*^*2*^ MB Nb clone, almost all the neurons produced were of γ type. **(F)** *dome*^*G0405*^ phenotype rescued by overexpression of *UAS-dome* in the clone. **(G)** *dome*^*G0405*^ phenotype rescued by overexpression of *UAS-Stat92EΔNΔC* in the clone. **(H)** 100% Stacked bar graph showing the penetrance of loss of JAK/STAT phenotypes and the rescue efficiency under overexpression of *dome* or *Stat92E*^*ΔNΔC*^. Genotype: (B) *FRT19A,UAS-mCD8-GFP/FRT19A,hs-FLP,tubP-GAL80; UAS-mCD8-GFP/+; GAL4-OK107/+*; (C) *FRT19A,dome*^*G0405*^*/FRT19A,hs-FLP,tubPGAL80; UAS-mCD8-GFP/+; GAL4-OK107/+*; (D) *FRT19A,dome*^*468*^*/FRT19A,hs-FLP,tubP-GAL80; UAS-mCD8-GFP/+; GAL4-OK107/+*; (E) *FRT19A,hop*^*2*^ */FRT19A,hs-FLP,tubP-GAL80; UAS-mCD8-GFP/+; GAL4-OK107/+*; (F) *FRT19A,dome*^*G0405*^*/FRT19A,hs-FLP,tubP-GAL80; UAS-mCD8-GFP/UAS-dome; GAL4-OK107/+*; (G) *FRT19A,dome*^*G0405*^*/FRT19A,hs-FLP,tubP-GAL80; UAS-mCD8-GFP/UAS-Stat92E* ^*ΔNΔC*^; *GAL4-OK107/+*. Scale bar, 25 μm.

### JAK/STAT pathway activity is not required for the survival and cell fate specification of MB neurons

Reduction in MB neurons could be due to decreased neuron generation or increased post-mitotic neuron death. Moreover, the γ-only phenotype of *dome* or *hop* MB could be the result of either cell fate specification failure or premature Nb termination. To clarify the cellular basis of *dome* or *ho*p MB phenotype, we first tested whether the JAK/STAT signaling affects post-mitotic neuron survival and cell fate specification of MB neurons. We generated *dome* MB Nb clones in NHL and examined them at different developmental stages (Fig. 2A-D). The numbers of neurons in the *dome* MB Nb clones were comparable (∼ 20) at newly emerged and 4-week-old adults (Fig. 2C,D), indicating that JAK/STAT is not required for the long-term survival of post-mitotic MB neurons. Furthermore, neurons generated by *dome* MB Nb clones were of normal γ neuron morphology at the wandering larval stage (Fig. 2A) and underwent remodeling at the pupal stage (Fig. 2B), which resulted in WT like adult γ neurons (Fig. 2C,D). We then generated *dome* MB Nb clones at different developmental stages and examined them in the adult brains (Fig. 2E-H). Whenever the clones were created, we invariably observed fewer neurons generated by *dome* MB Nb clones. However, the subtype switch of MB neurons wasn’t affected. For example, when created in the 2nd instar larvae (24 hrs ALH), *dome* MB Nb clones produced γ neurons and a few α’/β’ neurons (Fig. 2E). In comparison, when created in the early 3rd instar larvae (48 hrs ALH), *dome* MB Nb clones mainly produced α’/β’ and a few γ and α/β neurons (Fig. 2F). *dome* MB Nb clones induced at the middle 3rd instar (72 hrs ALH) produced α’/β’ and α/β neurons (Fig. 2G). And *dome* MB Nb clones induced at early pupa (96 hrs ALH) produced only α/β neurons (Fig. 2H). All together, these results suggest that JAK/STAT pathway does not affect the survival and cell fate specification of post-mitotic MB neurons.

**Figure 2.**
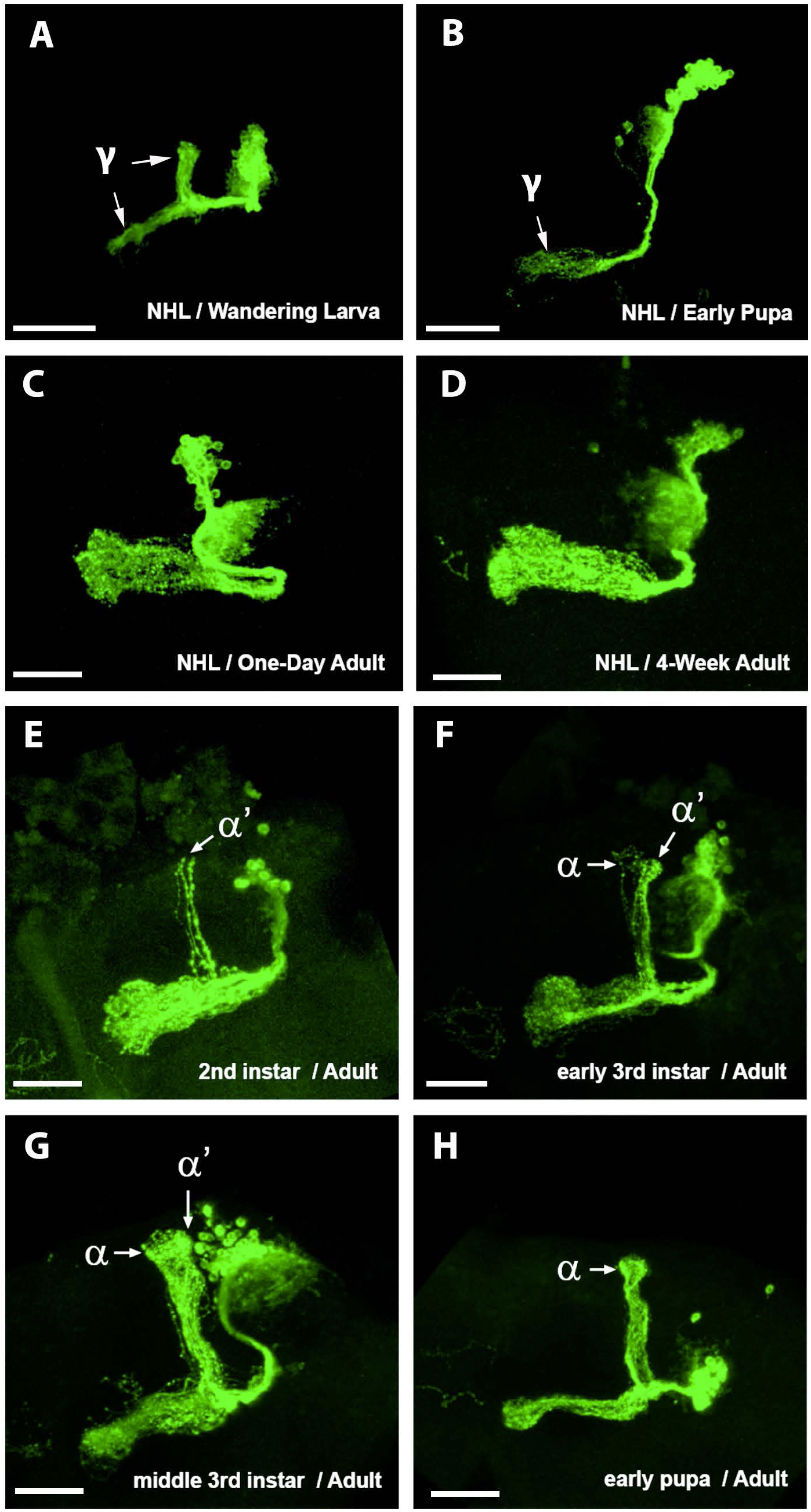
Loss of *dome* does not affect differentiation and survival of the postmitotic neurons. Representative confocal images of *dome* ^*G0405*^ MB Nb clones induced/examined at the indicated developmental stages. All clones were labeled by *GAL4-OK107>UAS-mCD8-GFP*. **(A-D)** *dome*^*G0405*^ MB Nb clone induced in the newly hatched larvae (NHL) and examined at different developmental stages, showing normal morphology **(A)** and remodeling **(B)** of larval γ neurons and the survival of γ adult neurons till 4 weeks after eclosion (comparing **D** to **C**). **(E-H)** *dome*^*G0405*^ MB Nb clone induced at different developmental stages and examined at one-week adults, showing normal switches of *dome* neurons from γ to α’/β’ and from α’/β’ to α/β. Consistent phenotypes were observed with > 5 clones for each panel. Genotype: *FRT19A,dome*^*G0405*^ */FRT19A,hs-FLP,tubPGAL80; UAS-mCD8-GFP/+; GAL4-OK107/+*. Scale bar, 25μm.

### JAK/STAT signaling prevents premature termination and promotes cell division of MB Nbs

To further clarify the cellular basis of this γ-only phenotype, we performed a time-course study to quantify neurons generated by wild type, *dome* loss-of-function, and Stat92E gain-of-function MB Nb clones. When created at NHL, each wild type MB Nb generated over 200 neurons. Note that this number is less than the reported ∼500 neurons generated by each MB Nb (Ito et al., 1997) (T. Lee et al., 1999). We reasoned this could be partially attributed to the way of counting cells. But it shouldn’t affect our conclusion, since we followed the consistent method for counting cells. In contrast to WT, the number of neurons generated by a *dome* mutant Nb stopped increasing at the early 3rd larval stage and remained at ∼20 hereafter. The number of neurons generated by a Stat92E gain-of-function Nb overpassed that generated by a WT Nb at all developmental stages and resulted in more than 500 neurons, a two-fold increase (Fig. 3A; Supplementary Fig. 1). We then labeled MB Nbs using an Nb-specific marker, Dpn antibody (Boone and Doe, 2008). At the early pupal stages, the brain hemisphere with a wild type clone had four MB Nbs (Fig. 3B). That with a *dome* mutant clone had only three MB Nbs all residing outside the clone (Fig. 3C), suggesting that the *dome* mutant MB Nb prematurely disappeared. However, the brain hemisphere with a Stat92E gain-of-function clone also had only four MB Nbs (Fig. 3D), indicating that the supernumerary neurons in Stat92E gain-of-function clones are probably a result of faster cell division from a single Nb. These data suggested that the JAK/STAT pathway might play two major roles in the MB neurogenesis: preventing Nb premature termination and promoting Nb cell proliferation.

**Figure 3.**
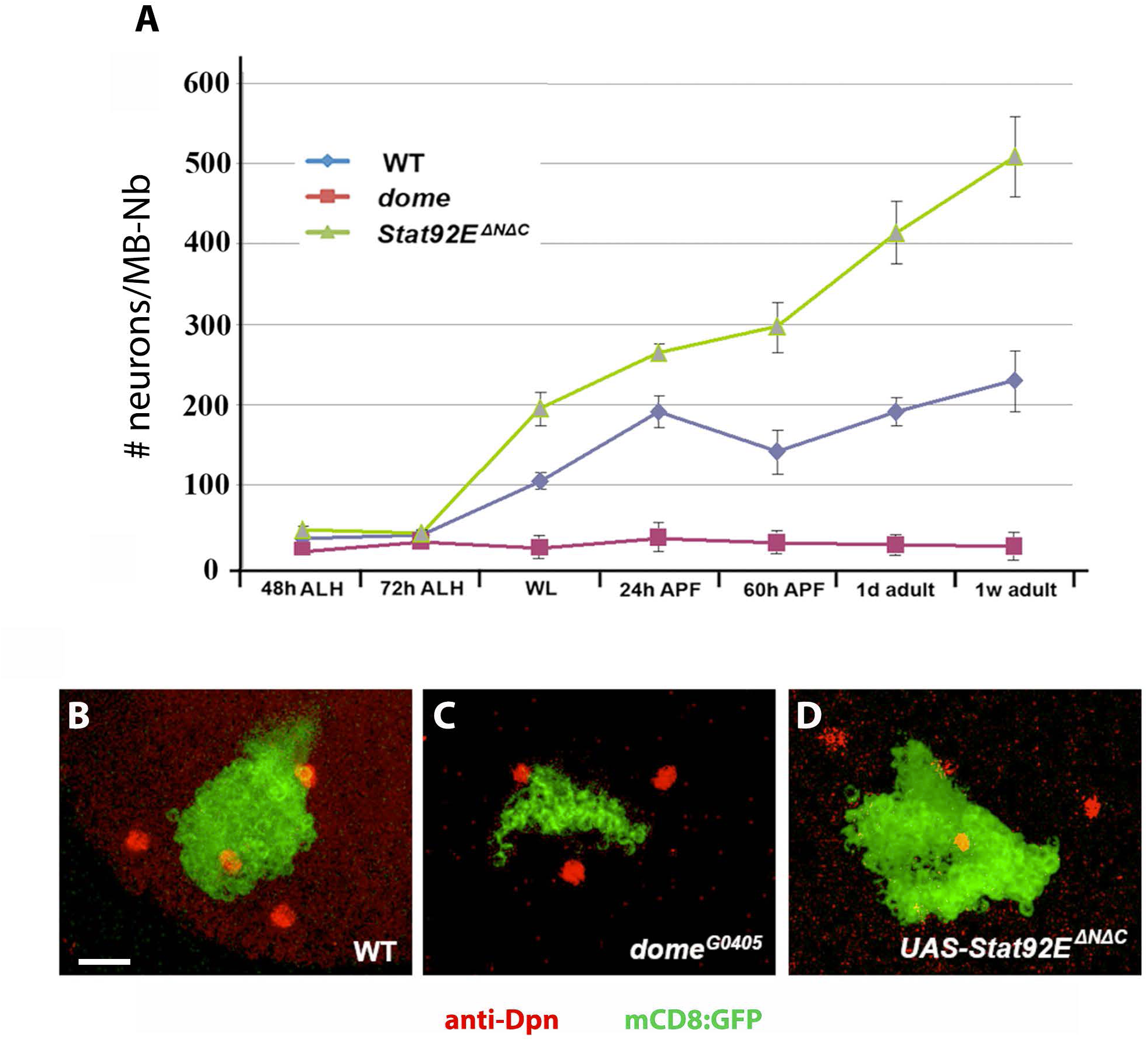
JAK/STAT signaling promotes division and prevents premature termination of MB Nbs. **(A)** Quantification of neurons generated by wild type (blue line), *dome* mutant (red line), or *Stat92E* gain-of-function (green line) MB Nb. MB Nb clones were induced in the newly hatched larvae and examined at different developmental stages. Data represent average of 12-30 clones. p < 0.0001 for WT vs *dome* and p < 0.01 for WT vs *Stat92E*^*ΔNΔC*^at all stages since WL; one-way ANOVA followed by Tukey’s honestly significant different (HSD) test was used to calculate statistical significance; see also Supplementary Figure 1. **(B-D)** Representative confocal images of the MB region of early pupal brains. Only cell bodies are presented to show MB Nb clones induced in the newly hatched larvae and labeled by *GAL4-OK107>UAS-mCD8-GFP* (green). Nbs were labeled by Dpn antibody staining (red). **(B)** WT; **(C)** *dome*^*G0405*^; and **(D)** expression of *UAS-Stat92E*^*ΔNΔC*^ in wild type clone. The image represents > 10 clones for each genotype. Genotype: (B) *FRT19A,UAS-mCD8-GFP/FRT19A,hs-FLP,tubP-GAL80; UAS-mCD8-FP/+; GAL4-OK107/+*; (C) *FRT19A,dome*|^*G0405*^*/FRT19A,hs-FLP,tubPGAL80; UAS-mCD8-GFP/+; GAL4-OK107/+*; (D) *FRT19A,mCD8-GFP/FRT19A,hs-FLP,tubP-GAL80; UAS-mCD8-GFP/UAS-Stat92E*^*ΔNΔC*^ *GAL4-OK107/+*. Scale bar, 25 μm.

### Overexpression of *CycE* and/or *diap1* significantly rescues loss of JAK/STAT pathway phenotypes

JAK/STAT signaling promoting cell proliferation in various tissues is well documented (Pellegrini and Dusanter-Fourt, 1997) (Levy and Darnell, 2002) (Arbouzova and Zeidler, 2006). However, its downstream targets that are required for this process are not clear (Zoranovic et al., 2013). To identify JAK/STAT signaling targets that are crucial for mediating MB neurogenesis, we collected UAS-transgene lines of 11 genes that are involved in the control of either cell proliferation, cell growth or cell death (Supplementary Table 1) and tested whether they could rescue the γ-only phenotype of *dome* MB Nb clones. Previous studies in mammals suggest *Cyclin D* (*CycD*) and *c-myc* as JAK/STAT downstream target genes to promote cell-cycle progression (Calò et al., 2003) (Bowman et al., 2000). We found that targeted overexpression of *CycD* and *myc* in the *dome* MB Nbs failed to rescue the mutant phenotype (Supplementary Fig. 2). Instead, overexpression of *CycE* in the *dome* MB Nb clones significantly rescued the γ-only phenotype, both α’/β’ and α/β neurons were restored based on the axon morphology (Fig. 4A,E). And overexpression of the *Drosophila inhibitor of apoptosis 1* (*diap1*) partially but significantly rescued the γ-only phenotype, with some α’/β’ neurons restored, but the later born α/β neurons were never found (Fig. 4B,E). Moreover, overexpression of *CycE* and *diap1* together resulted in more complete rescue than either of them separately, significant level of α’/β’ and α/β neurons were restored (Fig. 4C,E). The significant rescue effect by CycE (Fig. 4A,E) suggested that loss of JAK/STAT signaling activity mainly impaired cell cycle progression of MB Nbs. Notably, simultaneous blocking of cell death restored more complete MB neurogenesis (Fig. 4C,E), demonstrating that both impaired proliferation and premature cell death account for the observed defects in *dome* mutant MB Nb clones. Based on these results, we reasoned that JAK/STAT signaling might function through *diap1* to prevent Nb termination, because *diap1* is a known direct target of Stat92E (Betz et al., 2008). Further study is needed to verify that. On the other hand, JAK/STAT might function through *CycE* to promote Nb proliferation.

**Figure 4.**
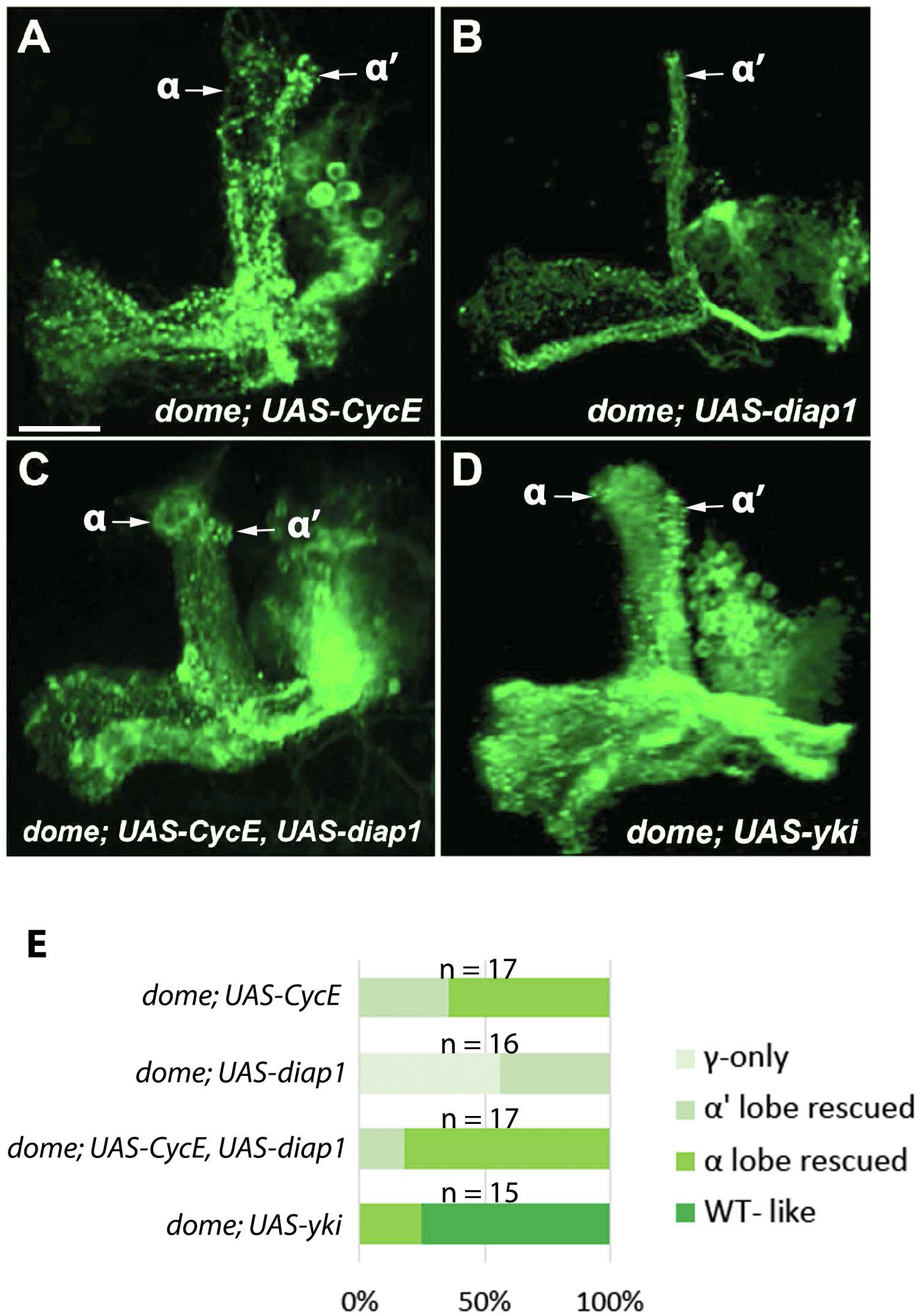
The neurogenesis defects of *dome* MB are rescued by *CycE, diap1*, or *yki*. Representative confocal images of *dome* MB Nb clones induced in the newly hatched larvae and examined in the adult brains, showing *dome*^*G0405*^ phenotypes are significantly rescued by excess *CycE***(A)** and partially rescued by excess *diap1***(B)**; substantially rescued by excess *CycE* and *diap1* together **(C)**; and fully rescued by excess *yki***(D). (E)** 100% Stacked bar graph showing the rescue efficiency of *dome* mutant phenotype under overexpression of *CycE* or/and *diap1*, or *yki*. Genotype: (A) *FRT19A,dome*^*G0405*^*/FRT19A,hs-FLP,tubP-GAL80; UAS-mCD8-GFP/+; UAS-CycE/+; GAL4-OK107/+*; (B) *FRT19A,dome*^*G0405*^*/FRT19A,hs-FLP,tubP-GAL80; UAS-mCD8-GFP/+; UAS-Diap1/+; GAL4-OK107/+*; (C) *FRT19A,dome*^*G0405*^*/FRT19A,hs-FLP,tubP-GAL80; UAS-mCD8-GFP/+; UAS-CycE,UAS-Diap1/+; GAL4-OK107/+*; (D) *FRT19A,dome*^*G0405*^*/FRT19A,hs-FLP,tubP-GAL80; UAS-mCD8-GFP/+; UAS-yki/+; GAL4-OK107/+*. Scale bar, 25 μm.

### Stat92E directly regulates *CycE* expression in MB Nbs to promote their proliferation

As overexpression of *CycE* substantially rescued the proliferation defects of *dome* Nbs (Fig. 4A,E), we were wondering whether *CycE* is a direct target that links JAK/STAT signaling to cell-cycle regulation. A consensus STAT-binding sequence, TTCNNNGAA (Horvath et al., 1995), was found in the regulatory region of *CycE* gene, which was also detected by whole genome ChIP-chip (modENCODE, Stat92E) (Supplementary Fig. 3A). This potential Stat92E-binding sequence is perfectly conserved across the 12 *Drosophila* genomes. To test whether it is a functional Stat92E response element in *vivo*, we produced a *lacZ* reporter transgenic fly line, *CycE(Stat92E-WT)-lacZ*, using an ∼ 1-kb genomic DNA fragment that contains this STAT-binding site (Supplementary Fig. 3B). *CycE(Stat92E-WT)-lacZ* expression was detected in the wing disc hinge, margin, and pouch regions (Supplementary Fig. 3D), which closely resembles the expression pattern of STAT-GFP, a Stat92E activity reporter (Bach et al., 2007). Moreover, when *Stat92E*^*ΔNΔc*^ was expressed in the posterior domain of wing disc driven by *en*-GAL4 (Neufeld et al., 1998), expression of *CycE(Stat92E-WT)-lacZ* increased in the corresponding region (Fig. 5A). To test whether the STAT-binding site in this fragment is responsible for the *lacZ* expression in wing disc, we generated another *lacZ* reporter transgenic fly line, *CycE(Stat92E-MT)-lacZ*, that carries a mutant STAT-binding site (TTCCAAGAA to TTCCAAGTT (Rivas et al., 2008)) (Supplementary Fig. 3A,B). *CycE(Stat92E-MT)-lacZ* was only weakly expressed in the wing disc (Supplementary Fig. 3E) and did not respond to Stat92E^ΔNΔC^ (Fig. 5B). These results suggested that Stat92E regulates *CycE* transcription via binding to the consensus STAT-binding site in *CycE* gene.

**Figure 5.**
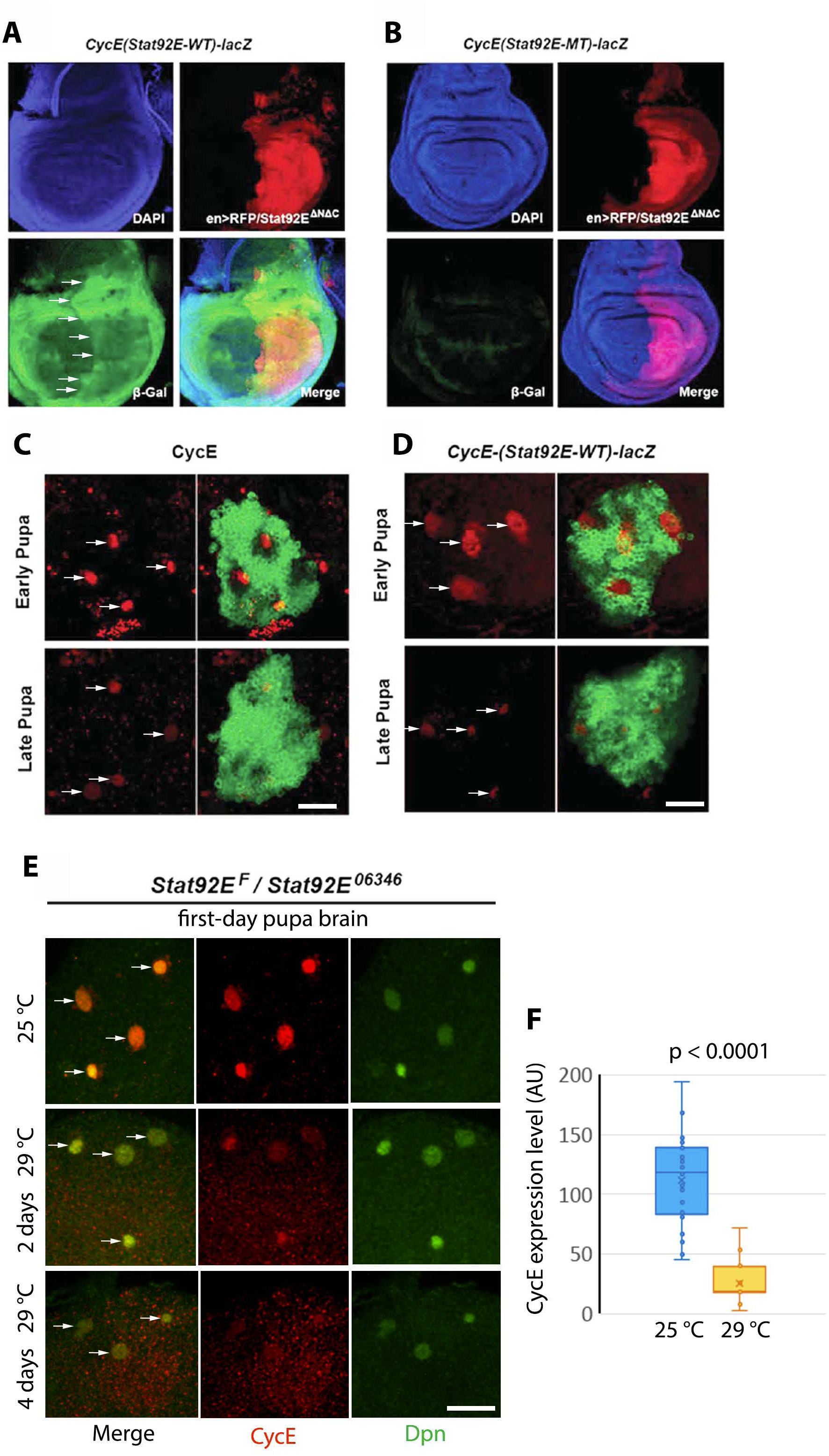
Stat92E directly regulates *CycE* transcription in MB Nbs and wing discs. **(A**,**B)** Representative confocal images of wing discs showing *CycE(Stat92E-WT)-lacZ* or *CycE(Stat92E-MT)-lacZ* expression (green, β-galactosidase antibody staining) in the *en-Gal4,UAS-Stat92E*^*ΔNΔC*^,*UAS-RFP* (red) genetic background. DAPI (blue) is used to label nuclei. Arrows, induced *CycE(Stat92E-WT)-lacZ* expression in the *en*-expressing region of the wing disc. **(C**,**D)** Representative confocal images showing cell body region of MB complex at pupal stages. CycE or *CycE(Stat92E-WT)-lacZ* expression (red, CycE or β-galactosidase antibody staining) was shown in the wild type MB labeled by GAL4-OK107>UAS-mCD8-GFP (green). Arrows, MB Nbs labeled with CycE or *CycE(Stat92E-WT)-lacZ*. **(E)** CycE (red, CycE antibody staining) and Dpn (green, Dpn antibody staining) levels in the *Stat92E* ^*ts*^ (*Stat92EF/Stat92E* ^*06346*^) MBs at the first day pupal stage. Temperature and days after switch are shown. Arrows, Nbs co-stained with Dpn and CycE. **(F)** Box plots comparing the CycE levels in MB Nbs of the first-day pupa brain from flies always incubated at 25°C (n = 32 Nbs from 8 biological samples) and flies switched to 29°C (n = 16 Nbs from 4 biological samples) for 2 days. Two-tailed t-test was used to calculate the statistical significance. Scale bar, 25 μm.

We then examined *CycE* and *CycE(Stat92E-WT)-lacZ* expression in MB and their dependence on the JAK/STAT pathway activity. Immunofluorescence staining with a CycE antibody revealed that CycE was broadly expressed in larval brains, and then mostly restricted to the area of four MB Nbs in each brain hemisphere during pupal stages. The CycE expression levels in MB Nb area were higher at early pupal stages and decreased at late pupal stages (Fig. 5C). This expression pattern correlates with the pattern of Nb proliferation during brain development – most Nbs generate neurons in the larval stages and terminate before pupal formation, but exceptionally, MB Nbs continuously divide until the end of pupal stage (Sousa-Nunes et al., 2010). *CycE(Stat92E-WT)-lacZ* showed a similar expression pattern to that of endogenous CycE, but was expressed in a broader area surrounding MB Nbs (Fig. 5D), which is, likely, because β-galactosidase is more stable than CycE protein. Also, it’s notable that *CycE(Stat92E-WT)-lacZ* showed highly restricted MB Nb-specific expression. In contrast, expression of *CycE(Stat92E-MT)-lacZ* was barely detected in the MB area of pupal brains (Supplementary Fig. 4), indicating that the predicted Stat92E-binding site is indispensable to the *CycE* expression in the MB Nbs.

To further test the dependence of *CycE* expression on Stat92E in MB Nbs, we performed a loss-of-function analysis using a temperature-sensitive Stat92E mutation, *Stat92E*^*F*^*/Stat92E*^*06346*^ (*Stat92E*^*ts*^) (Baksa et al., 2002). Dpn and CycE were co-stained to show the Nbs and the CycE expression levels. Compared to those consistently kept at permissive temperature (25°C), significantly decreased CycE levels were observed in the MB Nbs of first-day pupae 2-4 days after the *Stat92E*^*ts*^ larvae were shifted to the restrictive temperature (29°C) (Fig. 5E,F). It’s notable that with 4 days incubation at 29°C, CycE levels were almost not detectable in the MB of first-day pupa brain, and Dpn expression was also greatly reduced in those samples, indicating problems in Nb proliferation and maintenance. Therefore, JAK/STAT signaling directly regulates *CycE* expression levels in MB Nbs. Based on all these results, we reasoned that JAK/STAT functions through *CycE* to promote proliferation of MB Nbs.

Finally, we tested the loss-of-function phenotypes of *CycE* in MB. When *CycE*^*AR95*^ (a *CycE* loss of function allele) MB Nb clones were generated at NHL, exactly the same γ-only phenotype as in loss of JAK/STAT MB Nb clones was observed in most cases (Fig. 6A). This result confirms that CycE is required for MB neurogenesis. Next we analyzed whether excess *Diap1* or *Stat92E*^*ΔNΔC*^ could rescue the *CycE* mutant phenotype. We found that neither *Diap1* overexpression nor *Stat92E*^*ΔNΔC*^in *CycE*^*AR95*^ MB Nb clones could compensate for the proliferation defects caused by loss of *CycE* (Fig. 6B,C), indicating that *CycE* functions downstream of JAK/STAT signaling to promote MB Nb proliferation. We further tested the effect of *CycE* gain-of-function in MB neurogenesis. Wild type, *CycE* mutant, and *CycE* gain-of-function MB Nb clones were induced in NHL, and the number of neurons generated were counted at one-week adults. Loss of *CycE* resulted in significantly smaller mutant clones. On the other side, expression of excess *CycE* in the wild-type clones led to neuronal overgrowth. Note that excess *CycE* leads to neuronal overgrowth to the same extend as excess Stat92E does (Figs. 6D,3A). Collectively, these results indicate that *CycE* is the major, if not the only, downstream target of JAK/STAT pathway in controlling MB Nb cell proliferation.

**Figure 6.**
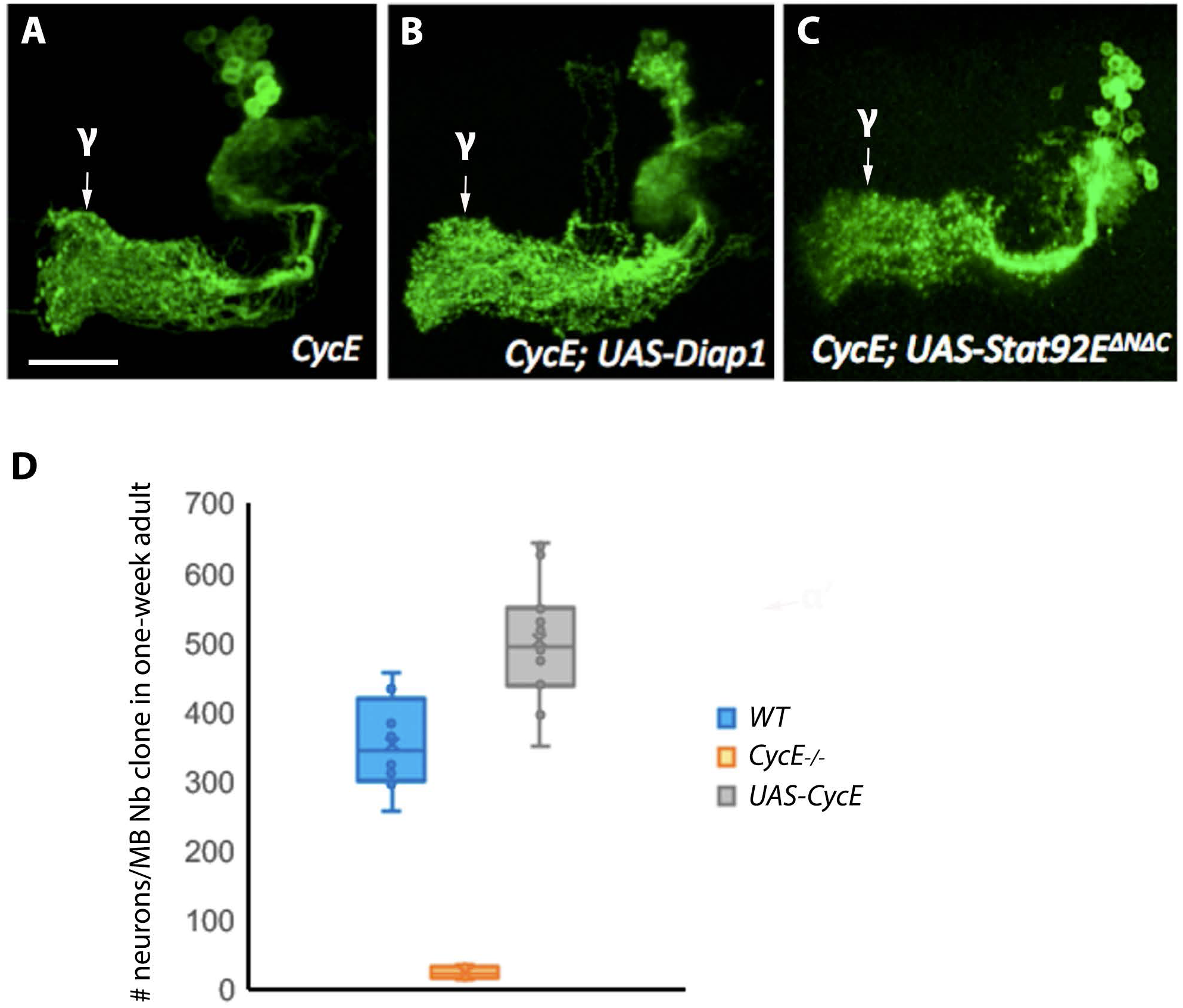
*CycE* mutant MB Nb clone phenocopies loss of JAK/STAT pathway activity, and excess *CycE* leads to neuronal overgrowth. **(A-C)** Representative confocal images of MARCM Nb clones induced in the newly hatched larvae and examined in the adult brains. MB neurons were labeled by *OK107-GAL4>UAS-mCD8-GFP*. **(A)** In *CycE*^*AR95*^ mutant clones, γ-only phenotype was observed. **(B)** In *CycE*^*AR95*^ clones with overexpression of *UAS-Diap1*, similar γ-only phenotype was observed. **(C)** In *CycE*^*AR95*^ clones with overexpression of *UAS-Stat92E*^*ΔNΔC*^, the clones also showed the γ-only phenotype. Consistent phenotypes were observed in > 8 samples for each genotype. **(D)** Box plot showing the number of neurons/MB Nb clone in one-week adult brains. The MARCM Nb clones were induced in NHL. Brains were collected in one-week adults and processed for mCD8 (clone marker) antibody staining. The number of neurons generated by wild type, *CycE* loss-of-function, and *CycE* gain-of-function MB Nbs was counted and compared. Loss of *CycE* results in significantly smaller mutant clones, with 24.5±2.6 cells (n=8 clones); expression of excess *CycE* in the wild-type clones leads to neuronal overgrowth, with 503±21.6 cells (n=16 clones), compared with 354±24.3 cells (n=8) in wild type clones. p < 0.01 for WT vs *CycE or UAS-CycE*; one-way ANOVA followed by Tukey’s honestly significant different (HSD) test was used to calculate statistical significance. Genotype: (A) *hs-FLP,UAS-mCD8-GFP/+; FRT40A,CycE*^*AR95*^*/FRTG40A,tubP-GAL80; GAL4-OK107/+*; (B) *hs-FLP,UAS-mCD8-GFP/+; FRT40A,CycE*^*AR95*^*/FRTG40A,tubPGAL80; UAS-Diap1/+; GAL4-OK107/+*; (C) *hs-FLP,UAS-mCD8-GFP/+; FRT40A,CycE*^*AR95*^*/FRTG40A,tubP-GAL80; UAS-Stat92E*^*ΔNΔC*^*/+; GAL4-OK107/+*; (D, WT) *hs-FLP,UAS-mCD8-GFP/+; FRTG13/FRTG13,tubP-GAL80; GAL4-OK107/+*; (D, *UAS-CycE*) *hs-FLP,UAS-mCD8-GFP/+; FRTG13/FRTG13,tubP-GAL80; UAS-CycE/+; GAL4-OK107/+*. Scale bar, 25 μm.

### JAK/STAT pathway functions in parallel with Hippo/Yorkie pathway in MB neurogenesis

While checking the rescue effect by various genes involved in cell growth, proliferation and cell death in loss of JAK/STAT MB Nb clones (Supplementary Table 1), we also found that overexpression of Hippo (Hpo) pathway downstream effector *yorkie* (*yki*) in the *dome* MB Nb clones fully rescued the γ-only phenotype of *dome* (Fig. 4D,E), which is consistent with previous reports that Yki is necessary and sufficient for NSC reactivation, growth and proliferation (Ding et al., 2016), and Hpo pathway restricts Nb proliferative potential and neuronal cell number through its inhibitory regulation on the key downstream transcription co-activator Yki (Poon et al., 2016). As *yki* overexpression fully rescues the loss of JAK/STAT phenotypes (Fig. 4D,E), it’s possible that Yki functions downstream to the JAK/STAT pathway or they function in parallel to control MB Nb proliferation. To test these hypotheses, we first checked the requirement of Yki function in normal MB development by generating MARCM clones of *yki*^*B5*^, an *yki* null allele (Bennett and Harvey, 2006), in MB. When induced at NHL, ∼43% *yki* Nb clones showed neurogenesis defects to different extents, such as reduced α’ or α lobes (Fig. 7A,B,D), suggesting premature Nb termination, while other clones had WT-like morphology (Fig. 7C). Thus, loss of *yki* in MB Nbs caused phenotypes similar to but less severe than loss of the JAK/STAT pathway activity. Then we tested whether higher JAK/STAT pathway activity could also rescue *yki* mutant phenotypes. When *Stat92E*^*ΔNΔC*^was expressed in *yki* MB Nb clones, *Stat92E*^*ΔNΔC*^fully rescued *yki* phenotypes (Fig. 7D). Therefore, loss of JAK/STAT pathway activity and loss of Hpo pathway downstream effector *yki* cause similar cell proliferation defects in MB, which can be mutually rescued by each other. These results suggested that JAK/STAT and Hippo/Yki pathways function in parallel to control MB Nb proliferation. Interestingly, *CycE* and *diap1* are also the major target genes of Yki, in regulating cell division and apoptosis (Tapon et al., 2002) (Wu et al., 2003) (Wu et al., 2008). To check whether Yki functions through *CycE* and/or *diap1* to regulate MB neurogenesis, we overexpressed *CycE* or/and *diap1* in *yki*^*B5*^ MB Nb clones. Overexpression of *CycE* and/or *diap1* partially but significantly rescued *yki* mutant phenotypes (Fig. 7D). Therefore, these results suggested that Yki might also function through *CycE* and *diap1* to sustain Nb proliferation. We propose that JAK/STAT and Yki coordinately control Nb proliferation by regulating the same downstream genes.

**Figure 7.**
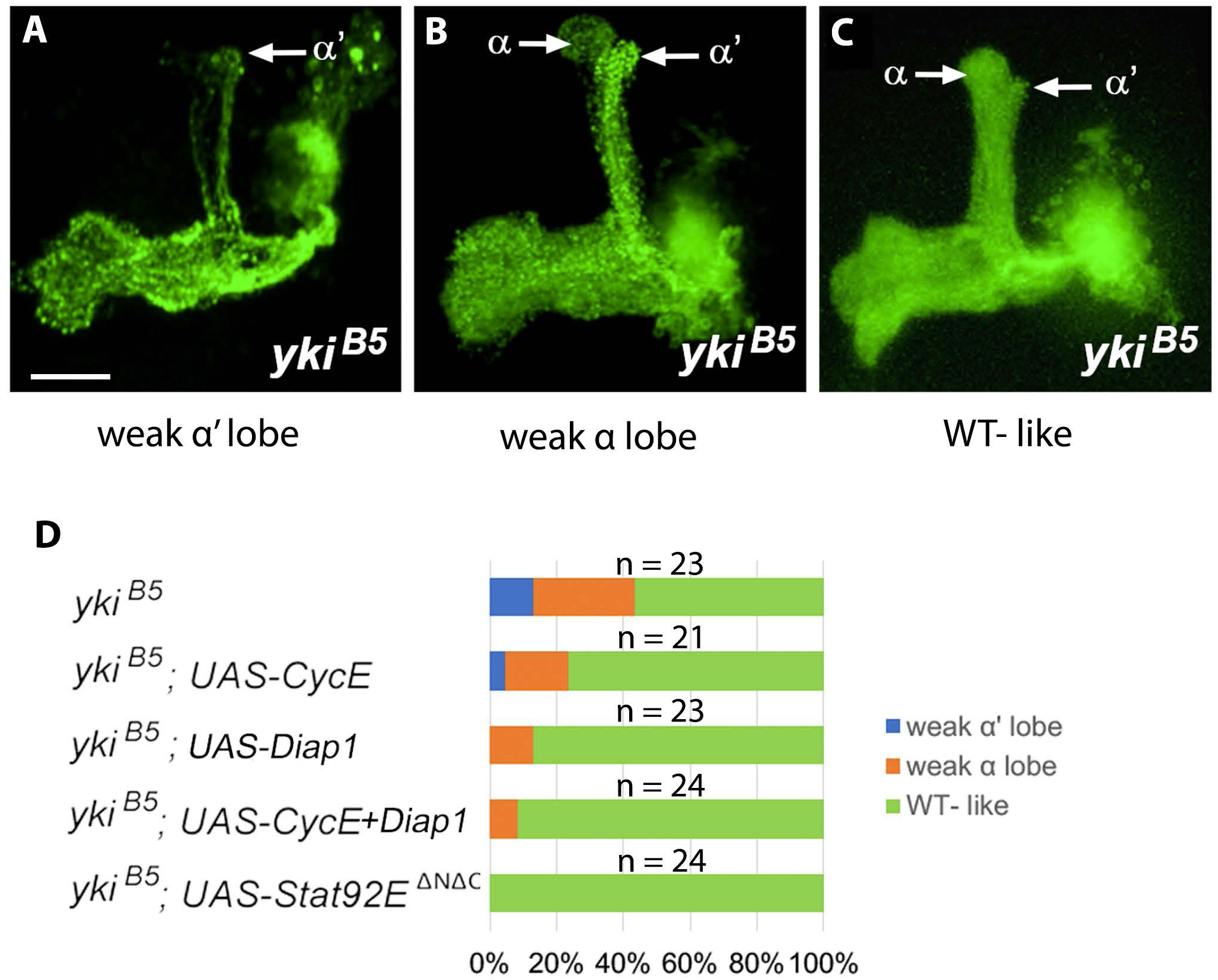
Loss of *yki* causes the similar but less severe defects as loss of JAK/STAT activity in MBs, which can be rescued by ectopic expression of *Stat92EΔNΔC* or *CycE* and/or *diap1*. **(A-C)** Representative confocal images of *yki*^*B5*^ Nb clones induced in newly hatched larvae and examined in adult brains. The lobe region is presented to show *yki*^*B5*^ phenotypes of different degrees. ∼43% *yki* Nb clones showed neurogenesis defects to different extends, such as reduced α’ or α lobes. **(A)** an example of weak α’ lobe; **(B)** an example of weak α lobe; **(C)** WT-like phenotype. **(D)** 100% Stacked bar graph showing the penetrance of *yki*^*B5*^ phenotypes of the reduced α or α’ lobes and the rescue efficiency under overexpression of *CycE, Diap1, CycE* plus *Diap1*, or *Stat92E*^*ΔNΔC*^. Genotype: (A-C) *hs-FLP,UAS-mCD8-GFP/+; FRTG13, yki*^*B5*^*/FRTG13,tubP-GAL80; GAL4-OK107/+*; (D, *yki*^*B5*^; *UAS-CycE*) *hs-FLP,UAS-mCD8-GFP/+; FRTG13,yki*^*B5*^*/FRTG13,tubPGAL80; UAS-CycE/+; GAL4-OK107/+*; (D, *yk*^*B5*^; *UAS-Diap1*) *hs-FLP,UAS-mCD8-GFP/+; FRTG13,yki*^*B5*^*/FRTG13,tubPGAL80; UAS-Diap1/+; GAL4-OK107/+*; (D, *yki*^*B5*^; *UAS-CycE+Diap1*) *hs-FLP,UAS-mCD8-GFP/+; FRTG13,yki*^*B5*^*/FRTG13,tubP-GAL80; UAS-CycE,UAS-Diap1/+; GAL4-OK107/+*; (D, *yki*^*B5*^; *UAS-Stat92E*^*ΔNΔC*^) *hs-FLP,UAS-mCD8-GFP/+; FRTG13,yki*^*B5*^*/FRTG13,tubP-GAL80; UAS-Stat92E*^*ΔNΔC*^*/+; GAL4-OK107/+*. Scale bar, 25 μm.

## Discussion

### *CycE* as the direct target of JAK/STAT to promote cell-cycle progression

Conserved roles for JAK/STAT signaling in the regulation of cell proliferation and stem cell maintenance have been reported in various metazoans (Pellegrini and Dusanter-Fourt, 1997) (Arbouzova and Zeidler, 2006). With only one receptor, Domeless (Dome), one JAK, Hopscotch (Hop), and one STAT, Stat92E, the lower complexity of *Drosophila* JAK/STAT pathway, together with the sophisticated genetic tools, has facilitated genetic dissection of this pathway. Two genes, *zfh1* and *chinmo*, were genetically identified as the Stat92E targets that mediate stem cell self-renewal (Leatherman and Dinardo, 2008) (Flaherty et al., 2010). However, the molecular downstream events by which the JAK/STAT pathway regulates cell proliferation are not yet clear (Zoranovic et al., 2013). Studies in the *Drosophila* eye and wing discs indicate that cells with excess JAK/STAT signaling activity progress through G1/S and G2/M cell cycle checkpoints faster than control disc cells (Bach et al., 2003) (Rodrigues et al., 2012), suggesting that JAK/STAT signaling directly or indirectly regulates the cell cycle progression factors, such as Cyclins and Cyclin-dependent kinases (Cdks). However, two independent genetic screens and three independent whole-genome RNAi screens did not identify a cell cycle gene as the downstream effector of JAK-STAT signaling pathway (Zoranovic et al., 2013). Here, we identify a solid connection between JAK/STST signaling and CycE, an essential G1 cyclin rate-limiting for progression into S phase, by evidencing that JAK/STAT controls Nb proliferation through the direct regulation of *CycE* transcription. In future, it will be important to check whether JAK/STAT pathway regulates cell proliferation through CycE in other tissues and morphogenetic contexts.

### Novel function of JAK/STAT in neurogenesis

The *Drosophila* central nervous system (CNS) develops from a bilateral neuroectoderm that contains a cluster of neuroepithelial (NE) cells. Selected NE cells differentiate into a few number of neuroblasts, which undergo asymmetric division producing a daughter neuroblast that self-renews, and a smaller ganglion mother cell (GMC) that gives rise to a variety of neuronal and glial cells. Previous studies in the optic lobe reveal that JAK/STAT pathway is required for NE cell maintenance and proliferation (W. Wang et al., 2011) and represses the transition of NE cells to Nbs (Yasugi et al., 2008) (Ngo et al., 2010). Here, we report that JAK/STAT pathway also plays important roles in neuroblast to promote its cell-cycle progression and prevent its premature termination.

Earlier studies have revealed several key regulators of MB Nb proliferation, including transcription factors Tailless (Tll) and Retinal homeobox (Rx) that positively regulate the mitotic activity and maintenance of MB Nbs (Kurusu et al., 2009) (Kraft et al., 2016), and Prospero (Pros) that is needed for termination of Nb cell proliferation (Li and Vaessin, 2000) (Choksi et al., 2006). It’s also shown that the levels of the temporal factors Imp and Syp exert important regulation on the Nb proliferative ability and timing for cell cycle exit (Liu et al., 2015) (Yang et al., 2017). And in MB, Imp and Syp control neuronal fates by regulating the temporal transcription factor Chinmo translation (Liu et al., 2015). A temporal gradient of Chinmo from high to low was shown to specify birth order-dependent neuronal fates (Zhu et al., 2006). Chinmo, on the other hand, has been identified as a key downstream effector of JAK/STAT signaling that mediate a variety of developmental and pathological processes including stem cell self-renewal (Flaherty et al., 2010). It will be interesting to see if JAK/STAT also functions through Chinmo to control MB Nb self-renewal and affect the temporal fates of MB neurons. Also, it will be important to know whether and how might JAK/STAT pathway interact with the other regulators of MB Nb proliferation discussed above.

### JAK/STAT and Hpo/Yki signaling pathways coordinately control MB Nb proliferation

The JAK/STAT and Hpo pathways are two major cell-proliferation-controlling signaling pathways in both vertebrates and invertebrates (Pellegrini and Dusanter-Fourt, 1997) (Arbouzova and Zeidler, 2006) (Pan, 2010). In *Drosophila*, they act through the prime effectors, Stat92E and Yki, respectively (Huang et al., 2005) (Hou et al., 1996) (Yan et al., 1996). Yki, a transcriptional coactivator, interacts with multiple DNA-binding transcription factors, such as Scalloped in *Drosophila* and its orthologs TEAD1-4 in vertebrates, to regulate the expression of the cell-cycle regulators *CycE* and the cell death inhibitor *diap1* (Wu et al., 2008) (Tapon et al., 2002). A previous study reports that JAK/STAT pathway directly regulates *diap1* expression (Betz et al., 2008). Here, we find that Stat92E directly regulates *CycE*. Therefore, JAK/STAT and Hpo signaling pathways coordinately control cell proliferation and cell death by regulating the same downstream genes. This notion is supported by the fact that loss of JAK/STAT or Yki activity causes the similar Nb proliferation defect, which can be mutually rescued by the overactivation of one another and by overexpression of *CycE* and *diap1* (Figs. 1,4,7). The function of Yki in promoting Nb proliferation is well known (Ding et al., 2016) (Poon et al., 2016). Our genetic evidence suggests that Yki functions through *CycE* and *diap1* to maintain Nb proliferative potential (Fig. 7D). These findings not only provide novel insights regarding the molecules that are downstream of the JAK/STAT or Yki in promoting Nb proliferation, but also shed light on the means by which distinct pathways converge to regulate the same biological process. Important future directions point to both the downstream and upstream regulations. Like on the downstream side, what are the *cis*-regulatory elements of *CycE* or *diap1* genes that are responsible for Stat92E and Yki regulation? Whereas on the upstream side, it’s interesting to know how JAK/STAT and Hpo/Yki signaling activities are tightly controlled for coordinated regulation of Nb proliferative activity.

## Materials and Methods

### Fly strains

Fly lines, including *dome*^*G0405*^, *dome*^*468>*^, *hop*^*2*^ and *CycE*^*AR95*^, were from Bloomington *Drosophila* Stock Center. *Stat92E*^*F*^, *Stat92E*^*06346*^, and *UAS-dome* were gifts from Steven Hou. *UAS-stat92E*^*ΔNΔC*^ was a gift from Erika Bach. UAS lines listed in the Supplementary Table 1, such as *UAS-CycE, UAS-diap1*, and *UAS-yorkie*, were gifts from Jianhua Huang. *yki*^*B5*^ was a gift from Duojia Pan. All flies were maintained on standard cornmeal medium at 25°C unless otherwise stated. *CycE(Stat92E-WT)-lacZ* and *CycE(Stat92E-MT)-lacZ* transgenic reporter lines were produced by P-element-mediated germline transformation. To make *CycE(Stat92E-WT)-lacZ* construct, ∼1kb DNA fragment flanking the predicted STAT-binding site in the *CycE* locus was amplified by genomic DNA PCR. The fragment was subcloned into the P-element transformation vector, pH-Pelican. To make *CycE(Stat92E-MT)-lacZ* construct, site-directed mutagenesis using Pfu DNA polymerase was performed with complementary oligonucleotides and DpnI digestion of the parental template. Products were transformed in DNA adenine methylation-free bacteria, tested for the successful generation of the TTCCAAGAA to TTCCAAGTT mutation by DNA sequencing.

### MARCM, immunohistochemistry and microscopy

MARCM clone induction in the brains was performed as described (T. Lee and Luo, 1999). To induce MARCM clones, *FRT19A,dome*^*G0405*^, *FRT19A,dome*^*468>*^, or *FRT19A,hop*^*2*^ flies were crossed to *FRT19A,hs-FLP,tubP-GAL80/Y; UAS-mCD8-GFP; GAL4-OK107* flies. *FRTG13,yki*^*B5*^ flies were crossed to *hs-FLP,UAS-mCD8-GFP; FRTG13,tubP-GAL80; GAL4-OK107* flies. And *FRT40A,CycE*^*AR95*^ flies were crossed to *hs-FLP,UAS-mCD8-GFP; FRTG40A,tubP-GAL80; GAL4-OK107* flies. Then F1 larvae were heat-shocked at 37°C for one hour at the indicted developmental stages. Brains were dissected out of larvae, pupae or adult flies with the right genotypes. Antibody staining was performed essentially as described (J. Wang et al., 2002). The following primary antibodies were used: rat anti-Dpn antibody (a gift from Chris Doe) (1: 10); rabbit anti-β-Galactosidase antibody (1: 50); and rabbit anti-CycE antibody (Santa Cruz Biotechnology) (1:50). FITC and Cy3 conjugated secondary antibodies were purchased from Jackson ImmunoResearch. All images of immunofluorescent staining were collected using a Zeiss LSM710 confocal microscope and processed with Adobe Photoshop.

### Time-course analysis

Wild type, *dome* loss-of-function, or *Stat92E* gain-of-function MB Nb clones were induced at newly hatched larvae. Brains were collected at 48h-, 72h-, 96h-ALH (after larvae hatching), 24h-, 60h-APF (after pupa formation), and 1day-, 1week-adults. Images of MB cell bodies were collected using confocal microscopy in 1 μm section. Individual Nb lineage sizes were determined by counting the number of cells through the optical sections in each clone.

### CycE expression analysis in *Stat92E* mutant brains

*Stat92E*^*F*^ flies were crossed with *Stat92E*^*06346*^*/TM3,GFP* flies at permissive temperature 25°C to generate two vials of synchronized progeny. When most larvae developed to early 3rd instar stages, one vial was kept at 25°C and another was moved to restrictive temperature 29°C. Two or four days after that, white pupae of *Stat92EF/Stat92E*^*06346*^ (*Stat92E*^*ts*^) were collected from both vials based on the absence of GFP. CycE expression in the brains was assessed by CycE antibody staining. Co-staining with Dpn antibody was done to label the Nbs. All the images were taken with the same confocal settings. CycE expression levels in Nbs were determined by measuring the fluorescence intensities with background subtracted.

### Statistical analyses

Statistical significance was determined by the two-tailed t-test (Fig. 5F), or one-way ANOVA (Figs. 3A,6D) followed by Tukey honestly significant different (HSD) test.

## Acknowledgements

We thank Erika Bach, Jianhua Huang, Steven Hou, Duojia Pan, Chris Doe, and the Bloomington Drosophila Stock Center for fly stocks and antibodies; Leslie Pick, Steven Hou, and Steve Mount for critical comments on the manuscript; Dr. AE Beaven for the UMD imaging core facility.

## Figure Legends

**Supplementary Figure 1**. Data for the quantification of neurons generated by wild type, *dome* mutant, or *Stat92E* gain-of-function MB Nb. MB Nb clones were induced in the newly hatched larvae. # cells per clone was quantitated at different developmental stages; (A) larval stages; (B) pupal stages; (C) adult stages.

**Supplementary Figure 2**. Excess *CycD*.*CDK4* or *myc* in the *dome* mutant MB Nbs failed to rescue the γ-only phenotype. Representative confocal images of *dome* MB Nb clones induced in the newly hatched larvae and examined in the adult brains, showing *UAS-CycD*.*CDK4***(A)** or *UAS-myc***(B)** failed to rescue *dome*^*G0405*^phenotypes.

**Supplementary Figure 3**. *CycE* gene structure and *CycE-lacZ* reporter transgenic lines. **(A)** The structure of *CycE* locus showing the consensus STAT-binding site (green; 2L: 15,744,093-15,744,102; TTCCAAGAA), which perfectly matches the STAT-binding site detected by ChIP-chip (red; 2L: 15,743,596-15744,299), and the position of different *lacZ* reporter transgenic lines. The *CycE-lacZ* reporter contains 16.4kb *CycE* cis-regulatory sequence including the consensus STAT-binding site and the *CycE(Stat92E-WT)-lacZ* and *CycE(Stat92E-MT)-lacZ* reporters contain a 1kb DNA segment that contains the predicted *CycE* STAT DNA-binding site. **(B)** Constructs for *CycE(Stat92E-WT)-lacZ* and *CycE(Stat92E-MT)-lacZ* reporter transgenic lines. The predicted *CycE* STAT DNA-binding site was mutated from TTCCAAGAA to TTCCAAGTT in *CycE(Stat92E-MT)-lacZ* construct. **(C)** A diagram of *Drosophila* wing imaginal disc modified from (Butler et al., 2003). **(D**,**E)** Expression of *CycE(Stat92E-WT)-lacZ* and *CycE(Stat92E-MT)-lacZ* in wing discs assessed by β-galactosidase antibody staining, showing that *CycE(Stat92E-WT)-lacZ* highly and *CycE(Stat92E-MT)-lacZ* weakly expresses in the hinge and pouch of wing discs.

**Supplementary Figure 4**. *CycE(Stat92E-MT)-lacZ* expression is not detectable in MB Nbs. Representative confocal images showing MB cell body region of GAL4-OK107>UAS-mCD8-GFP; *CycE(Stat92E-MT)-lacZ* at early pupal stage. Green: mCD8-GFP, labeling MB neurons. Red: *CycE(Stat92E-MT)-lacZ* expression detected by β-galactosidase antibody staining. Scale bar, 25 μm.

**Supplementary Table 1**. The list of UAS-transgene lines of 11 genes involved in the control of either cell proliferation, cell growth or cell death. Each line was used to test the rescue efficiency of the *dome* mutant MB Nb clones. The rescue efficiency was concluded based on at least 15 clones each genotype.

## References

Arbouzova, N.I., Zeidler, M.P., 2006. JAK/STAT signalling in Drosophila: insights into conserved regulatory and cellular functions. Development (Cambridge, England) 133, 2605–2616. doi: 10.1242/dev.02411

Bach, E.A., Ekas, L.A., Ayala-Camargo, A., Flaherty, M.S., Lee, H., Perrimon, N., Baeg, G.-H., 2007. GFP reporters detect the activation of the Drosophila JAK/STAT pathway in vivo. Gene Expr. Patterns 7, 323–331. doi: 10.1016/j.modgep.2006.08.003

Bach, E.A., Vincent, S., Zeidler, M.P., Perrimon, N., 2003. A sensitized genetic screen to identify novel regulators and components of the Drosophila janus kinase/signal transducer and activator of transcription pathway. Genetics 165, 1149–1166.

Baksa, K., Parke, T., Dobens, L.L., Dearolf, C.R., 2002. The Drosophila STAT protein, stat92E, regulates follicle cell differentiation during oogenesis. Developmental biology 243, 166–175. doi: 10.1006/dbio.2001.0539

Bello, B., Reichert, H., Hirth, F., 2006. The brain tumor gene negatively regulates neural progenitor cell proliferation in the larval central brain of Drosophila. Development (Cambridge, England) 133, 2639–2648. doi: 10.1242/dev.02429

Bennett, F.C., Harvey, K.F., 2006. Fat cadherin modulates organ size in Drosophila via the Salvador/Warts/Hippo signaling pathway. Current biology: CB 16, 2101–2110. doi: 10.1016/j.cub.2006.09.045

Betschinger, Joerg, Mechtler, K., Knoblich, J.A., 2006. Asymmetric segregation of the tumor suppressor brat regulates self-renewal in Drosophila neural stem cells. Cell 124, 1241–1253. doi: 10.1016/j.cell.2006.01.038

Betschinger, Jörg, Knoblich, Jürgen A, 2004. Dare to be different: asymmetric cell division in Drosophila, C. elegans and vertebrates. Current biology: CB 14, R674–85. doi: 10.1016/j.cub.2004.08.017

Betz, A., Ryoo, H.D., Steller, H., Darnell, J.E., 2008. STAT92E is a positive regulator of Drosophila inhibitor of apoptosis 1 (DIAP/1) and protects against radiation-induced apoptosis. Proceedings of the National Academy of Sciences of the United States of America 105, 13805–13810. doi: 10.1073/pnas.0806291105

Boone, J.Q., Doe, C.Q., 2008. Identification of Drosophila type II neuroblast lineages containing transit amplifying ganglion mother cells. Dev Neurobiol 68, 1185–1195. doi: 10.1002/dneu.20648

Bowman, T., Garcia, R., Turkson, J., Jove, R., 2000. STATs in oncogenesis. Oncogene 19, 2474–2488. doi: 10.1038/sj.onc.1203527

Butler, M.J., Jacobsen, T.L., Cain, D.M., Jarman, M.G., Hubank, M., Whittle, J.R.S., Phillips, R., Simcox, A., 2003. Discovery of genes with highly restricted expression patterns in the Drosophila wing disc using DNA oligonucleotide microarrays. Development (Cambridge, England) 130, 659–670. doi: 10.1242/dev.00293

Calò, V., Migliavacca, M., Bazan, V., Macaluso, M., Buscemi, M., Gebbia, N., Russo, A., 2003. STAT proteins: from normal control of cellular events to tumorigenesis. J. Cell. Physiol. 197, 157–168. doi: 10.1002/jcp.10364

Choksi, S.P., Southall, T.D., Bossing, T., Edoff, K., de Wit, E., Fischer, B.E., van Steensel, B., Micklem, G., Brand, A.H., 2006. Prospero acts as a binary switch between self-renewal and differentiation in Drosophila neural stem cells. Developmental cell 11, 775–789. doi: 10.1016/j.devcel.2006.09.015

Ding, R., Weynans, K., Bossing, T., Barros, C.S., Berger, C., 2016. The Hippo signalling pathway maintains quiescence in Drosophila neural stem cells. Nature communications 7, 10510–12. doi: 10.1038/ncomms10510

Doe, C.Q., 2008. Neural stem cells: balancing self-renewal with differentiation. Development (Cambridge, England) 135, 1575–1587. doi: 10.1242/dev.014977

Ekas, L.A., Cardozo, T.J., Flaherty, M.S., McMillan, E.A., Gonsalves, F.C., Bach, E.A., 2010. Characterization of a dominant-active STAT that promotes tumorigenesis in Drosophila. Developmental biology 344, 621–636. doi: 10.1016/j.ydbio.2010.05.497

Flaherty, M.S., Salis, P., Evans, C.J., Ekas, L.A., Marouf, A., Zavadil, J., Banerjee, U., Bach, E.A., 2010. chinmo is a functional effector of the JAK/STAT pathway that regulates eye development, tumor formation, and stem cell self-renewal in Drosophila. Developmental cell 18, 556–568. doi: 10.1016/j.devcel.2010.02.006

Gage, F.H., Temple, S., 2013. Neural stem cells: generating and regenerating the brain. Neuron 80, 588–601. doi: 10.1016/j.neuron.2013.10.037

Gómez-López, S., Lerner, R.G., Petritsch, C., 2014. Asymmetric cell division of stem and progenitor cells during homeostasis and cancer. Cell. Mol. Life Sci. 71, 575–597. doi: 10.1007/s00018-013-1386-1

Homem, C.C.F., Knoblich, Juergen A, 2012. Drosophila neuroblasts: a model for stem cell biology. Development (Cambridge, England) 139, 4297–4310. doi: 10.1242/dev.080515

Horvath, P., Suganuma, A., Inaba, M., Pan, Y.B., Gupta, K.C., 1995. Multiple elements in the 5’ untranslated region down-regulate c-sis messenger RNA translation. Cell Growth Differ. 6, 1103–1110.

Hou, X.S., Melnick, M.B., Perrimon, N., 1996. Marelle acts downstream of the Drosophila HOP/JAK kinase and encodes a protein similar to the mammalian STATs. Cell 84, 411–419. doi: 10.1016/s0092-8674(00)81286-6

Huang, J., Wu, S., Barrera, J., Matthews, K., Pan, D., 2005. The Hippo signaling pathway coordinately regulates cell proliferation and apoptosis by inactivating Yorkie, the Drosophila Homolog of YAP. Cell 122, 421–434. doi: 10.1016/j.cell.2005.06.007

Ito, K., Awano, W., Suzuki, K., Hiromi, Y., Yamamoto, D., 1997. The Drosophila mushroom body is a quadruple structure of clonal units each of which contains a virtually identical set of neurones and glial cells. Development (Cambridge, England) 124, 761–771.

Kraft, K.F., Massey, E.M., Kolb, D., Walldorf, U., Urbach, R., 2016. Retinal homeobox promotes cell growth, proliferation and survival of mushroom body neuroblasts in the Drosophila brain. Mech. Dev. 142, 50–61. doi: 10.1016/j.mod.2016.07.003

Kurusu, M., Maruyama, Y., Adachi, Y., Okabe, M., Suzuki, E., Furukubo-Tokunaga, K., 2009. A conserved nuclear receptor, Tailless, is required for efficient proliferation and prolonged maintenance of mushroom body progenitors in the Drosophila brain. Developmental biology 326, 224–236. doi: 10.1016/j.ydbio.2008.11.013

Leatherman, J.L., Dinardo, S., 2008. Zfh-1 controls somatic stem cell self-renewal in the Drosophila testis and nonautonomously influences germline stem cell self-renewal. Cell Stem Cell 3, 44–54. doi: 10.1016/j.stem.2008.05.001

Lee, C.-Y., Robinson, K.J., Doe, C.Q., 2006a. Lgl, Pins and aPKC regulate neuroblast self-renewal versus differentiation. Nature 439, 594–598. doi: 10.1038/nature04299

Lee, C.-Y., Wilkinson, B.D., Siegrist, S.E., Wharton, R.P., Doe, C.Q., 2006b. Brat is a Miranda cargo protein that promotes neuronal differentiation and inhibits neuroblast self-renewal. Developmental cell 10, 441–449. doi: 10.1016/j.devcel.2006.01.017

Lee, T., Lee, A., Luo, L., 1999. Development of the Drosophila mushroom bodies: sequential generation of three distinct types of neurons from a neuroblast. Development (Cambridge, England) 126, 4065–4076.

Lee, T., Luo, L., 1999. Mosaic analysis with a repressible cell marker for studies of gene function in neuronal morphogenesis. Neuron 22, 451–461. doi: 10.1016/s0896-6273(00)80701-1

Levy, D.E., Darnell, J.E., 2002. Stats: transcriptional control and biological impact. Nat. Rev. Mol. Cell Biol. 3, 651–662. doi: 10.1038/nrm909

Li, L., Vaessin, H., 2000. Pan-neural Prospero terminates cell proliferation during Drosophila neurogenesis. Genes Dev. 14, 147–151.

Liu, Z., Yang, C.-P., Sugino, K., Fu, C.-C., Liu, L.-Y., Yao, X., Lee, L.P., Lee, T., 2015. Opposing intrinsic temporal gradients guide neural stem cell production of varied neuronal fates. Science 350, 317–320. doi: 10.1126/science.aad1886

Neufeld, T.P., la Cruz, de, A.F., Johnston, L.A., Edgar, B.A., 1998. Coordination of growth and cell division in the Drosophila wing. Cell 93, 1183–1193. doi: 10.1016/s0092-8674(00)81462-2

Ngo, K.T., Wang, J., Junker, M., Kriz, S., Vo, G., Asem, B., Olson, J.M., Banerjee, U., Hartenstein, V., 2010. Concomitant requirement for Notch and Jak/Stat signaling during neuro-epithelial differentiation in the Drosophila optic lobe. Developmental biology 346, 284–295. doi: 10.1016/j.ydbio.2010.07.036

Pan, D., 2010. The hippo signaling pathway in development and cancer. Developmental cell 19, 491–505. doi: 10.1016/j.devcel.2010.09.011

Pellegrini, S., Dusanter-Fourt, I., 1997. The structure, regulation and function of the Janus kinases (JAKs) and the signal transducers and activators of transcription (STATs). Eur. J. Biochem. 248, 615–633. doi: 10.1111/j.1432-1033.1997.00615.x

Poon, C.L.C., Mitchell, K.A., Kondo, S., Cheng, L.Y., Harvey, K.F., 2016. The Hippo Pathway Regulates Neuroblasts and Brain Size in Drosophila melanogaster. Current biology: CB 26, 1034–1042. doi: 10.1016/j.cub.2016.02.009

Resnitzky, D., Gossen, M., Bujard, H., Reed, S.I., 1994. Acceleration of the G1/S phase transition by expression of cyclins D1 and E with an inducible system. Mol. Cell. Biol. 14, 1669–1679. doi: 10.1128/mcb.14.3.1669

Rivas, M.L., Cobreros, L., Zeidler, M.P., Hombría, J.C.-G., 2008. Plasticity of Drosophila Stat DNA binding shows an evolutionary basis for Stat transcription factor preferences. EMBO reports 9, 1114–1120. doi: 10.1038/embor.2008.170

Rodrigues, A.B., Zoranovic, T., Ayala-Camargo, A., Grewal, S., Reyes-Robles, T., Krasny, M., Wu, D.C., Johnston, L.A., Bach, E.A., 2012. Activated STAT regulates growth and induces competitive interactions independently of Myc, Yorkie, Wingless and ribosome biogenesis. Development (Cambridge, England) 139, 4051–4061. doi: 10.1242/dev.076760

Sousa-Nunes, R., Cheng, L.Y., Gould, A.P., 2010. Regulating neural proliferation in the Drosophila CNS. Curr. Opin. Neurobiol. 20, 50–57. doi: 10.1016/j.conb.2009.12.005

Tapon, N., Harvey, K.F., Bell, D.W., Wahrer, D.C.R., Schiripo, T.A., Haber, D.A., Hariharan, I.K., 2002. salvador Promotes both cell cycle exit and apoptosis in Drosophila and is mutated in human cancer cell lines. Cell 110, 467–478. doi: 10.1016/s0092-8674(02)00824-3

Wang, J., Lee, C.-H.J., Lin, S., Lee, T., 2006. Steroid hormone-dependent transformation of polyhomeotic mutant neurons in the Drosophila brain. Development (Cambridge, England) 133, 1231–1240. doi: 10.1242/dev.02299

Wang, J., Zugates, C.T., Liang, I.H., Lee, C.-H.J., Lee, T., 2002. Drosophila Dscam is required for divergent segregation of sister branches and suppresses ectopic bifurcation of axons. Neuron 33, 559–571. doi: 10.1016/s0896-6273(02)00570-6

Wang, W., Li, Y., Zhou, L., Yue, H., Luo, H., 2011. Role of JAK/STAT signaling in neuroepithelial stem cell maintenance and proliferation in the Drosophila optic lobe. Biochem. Biophys. Res. Commun. 410, 714–720. doi: 10.1016/j.bbrc.2011.05.119

Wu, S., Huang, J., Dong, J., Pan, D., 2003. hippo encodes a Ste-20 family protein kinase that restricts cell proliferation and promotes apoptosis in conjunction with salvador and warts. Cell 114, 445–456. doi: 10.1016/s0092-8674(03)00549-x

Wu, S., Liu, Y., Zheng, Y., Dong, J., Pan, D., 2008. The TEAD/TEF family protein Scalloped mediates transcriptional output of the Hippo growth-regulatory pathway. Developmental cell 14, 388–398. doi: 10.1016/j.devcel.2008.01.007

Yan, R., Small, S., Desplan, C., Dearolf, C.R., Darnell, J.E., 1996. Identification of a Stat gene that functions in Drosophila development. Cell 84, 421–430. doi: 10.1016/s0092-8674(00)81287-8

Yang, C.-P., Samuels, T.J., Huang, Y., Yang, L., Ish-Horowicz, D., Davis, I., Lee, T., 2017. Imp and Syp RNA-binding proteins govern decommissioning of Drosophila neural stem cells. Development (Cambridge, England) 144, 3454–3464. doi: 10.1242/dev.149500

Yasugi, T., Umetsu, D., Murakami, S., Sato, M., Tabata, T., 2008. Drosophila optic lobe neuroblasts triggered by a wave of proneural gene expression that is negatively regulated by JAK/STAT. Development (Cambridge, England) 135, 1471–1480. doi: 10.1242/dev.019117

Yu, F., Kuo, C.T., Jan, Y.N., 2006. Drosophila neuroblast asymmetric cell division: recent advances and implications for stem cell biology. Neuron 51, 13–20. doi: 10.1016/j.neuron.2006.06.016

Zheng, X., Wang, J., Haerry, T.E., Wu, A.Y.-H., Martin, J., O’Connor, M.B., Lee, C.-H.J., Lee, T., 2003. TGF-beta signaling activates steroid hormone receptor expression during neuronal remodeling in the Drosophila brain. Cell 112, 303–315. doi: 10.1016/s0092-8674(03)00072-2

Zhu, S., Lin, S., Kao, C.-F., Awasaki, T., Chiang, A.-S., Lee, T., 2006. Gradients of the Drosophila Chinmo BTB-zinc finger protein govern neuronal temporal identity. Cell 127, 409–422. doi: 10.1016/j.cell.2006.08.045

Zoranovic, T., Grmai, L., Bach, E.A., 2013. Regulation of proliferation, cell competition, and cellular growth by the Drosophila JAK-STAT pathway. JAKSTAT 2, e25408. doi: 10.4161/jkst.25408

