## Supplementary figures and images for "JAK/STAT pathway promotes *Drosophila* neuroblast proliferation via the direct *CycE* regulation"

### Supplementary Figure 1

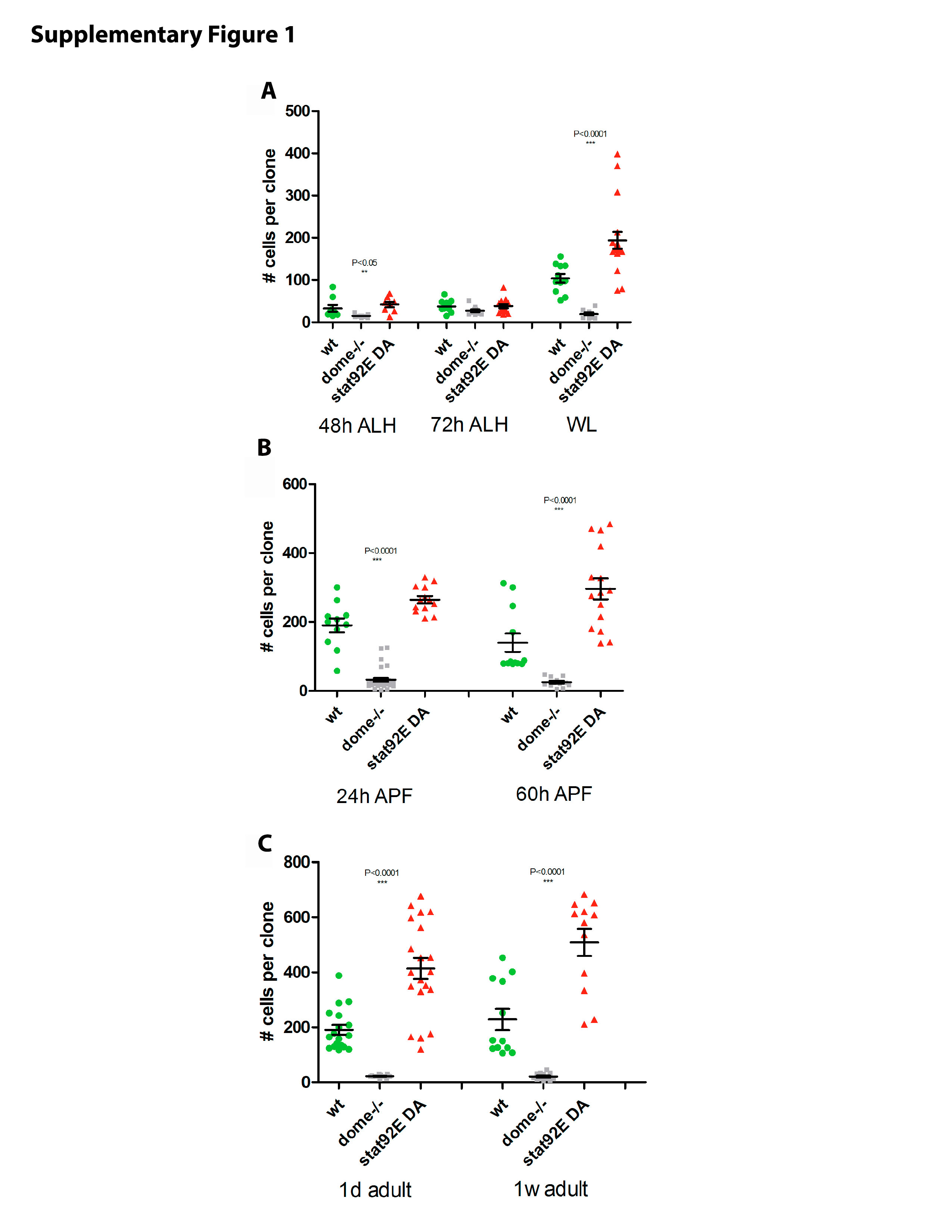

### Supplementary Figure 2

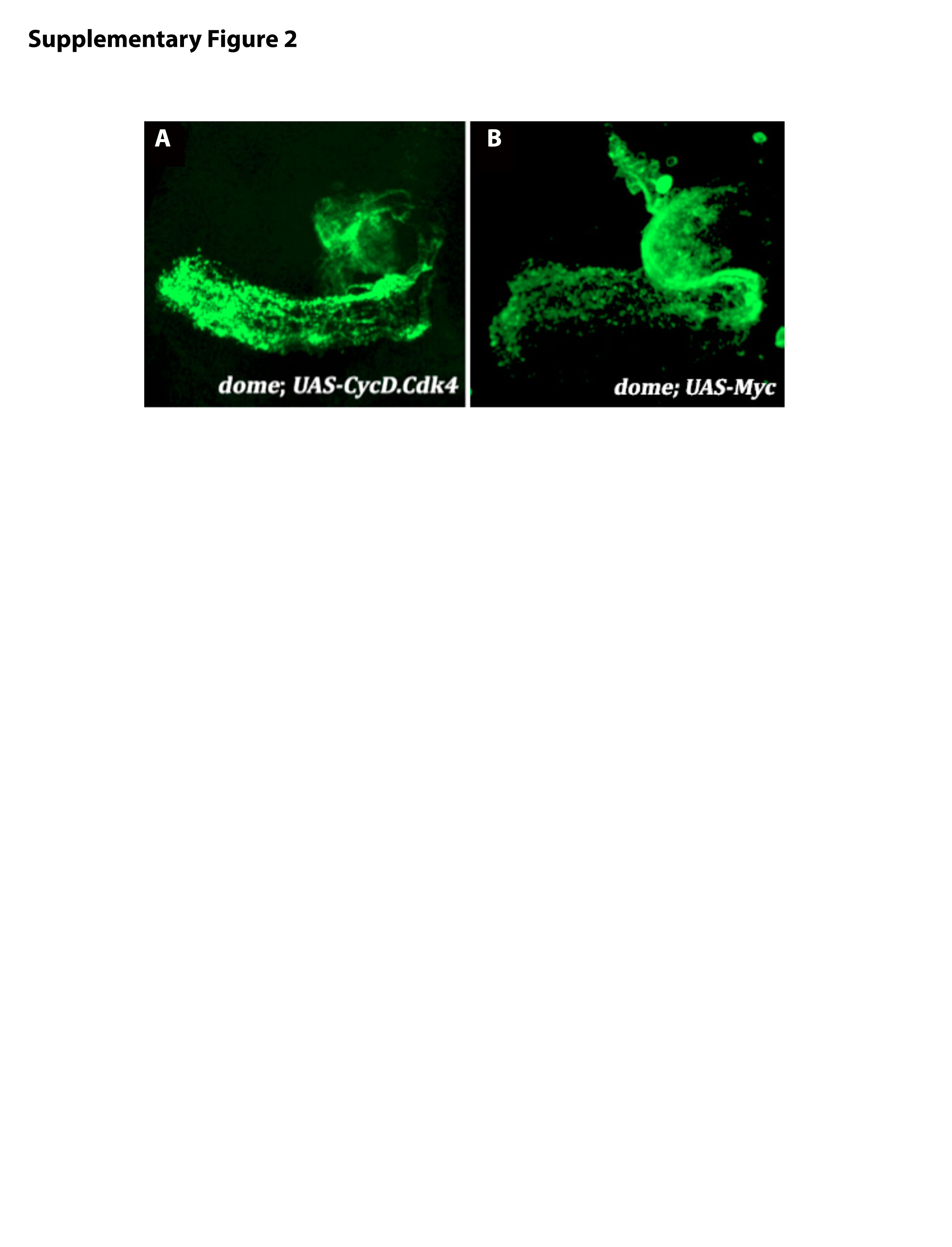

### Supplementary Figure 3

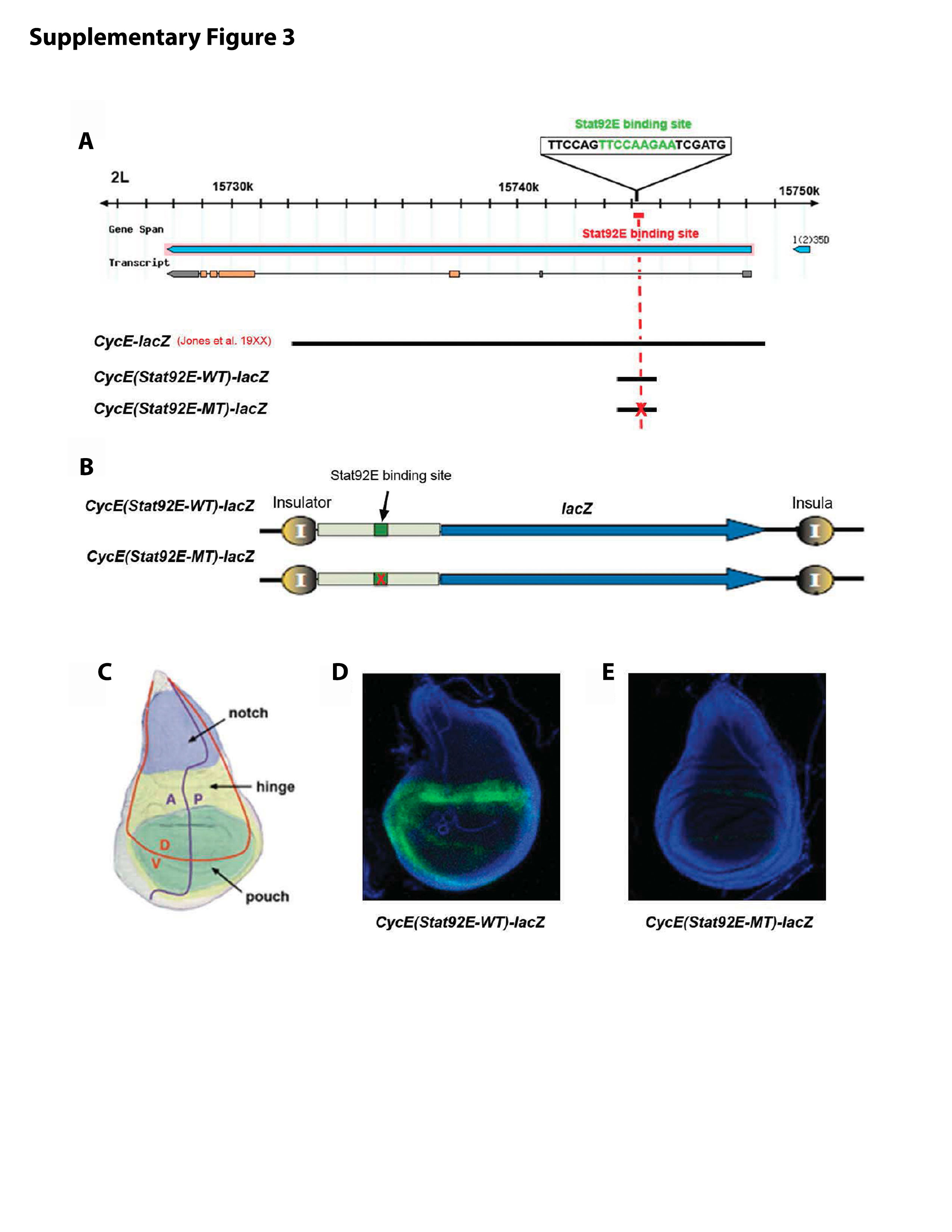

### Supplementary Figure 4

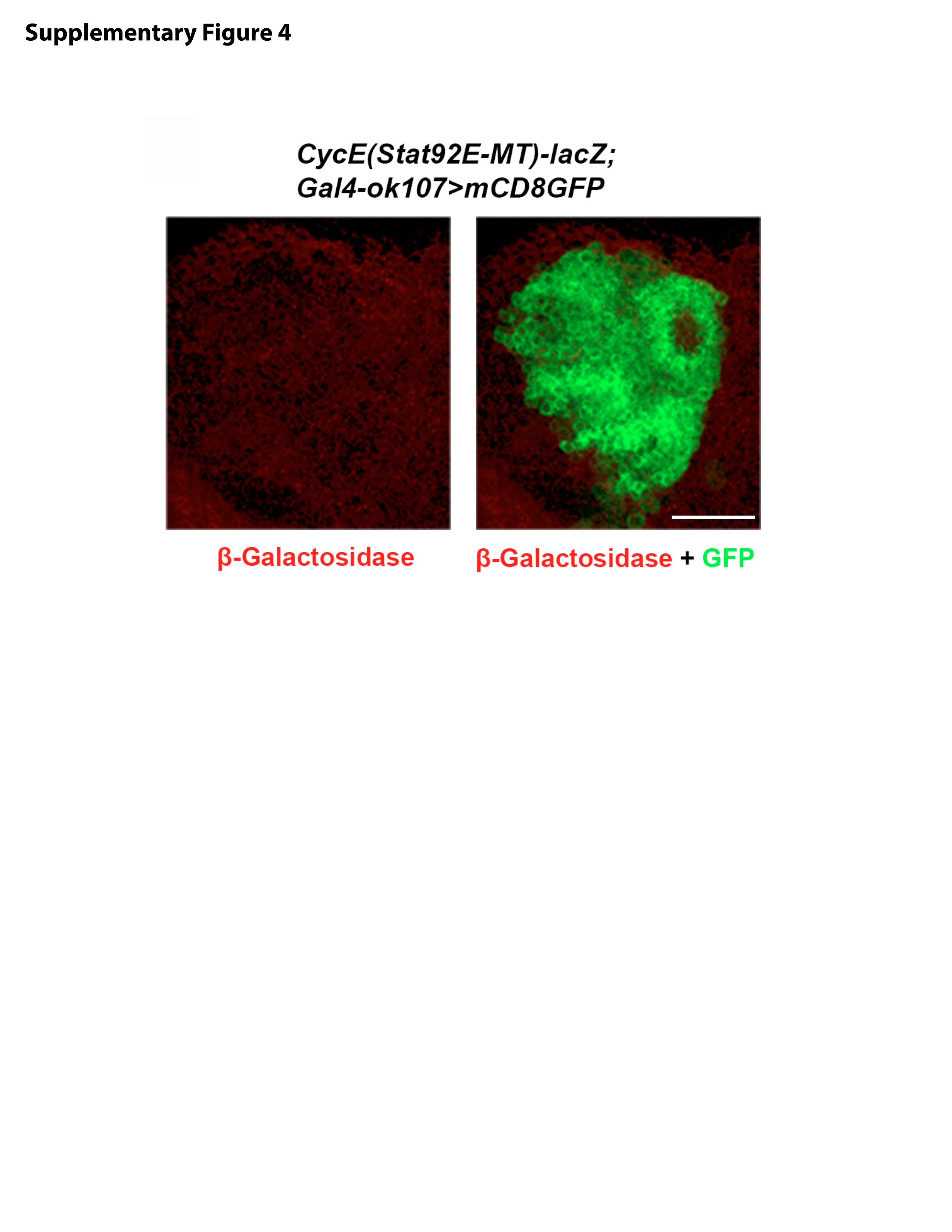

### Supplementary Table 1

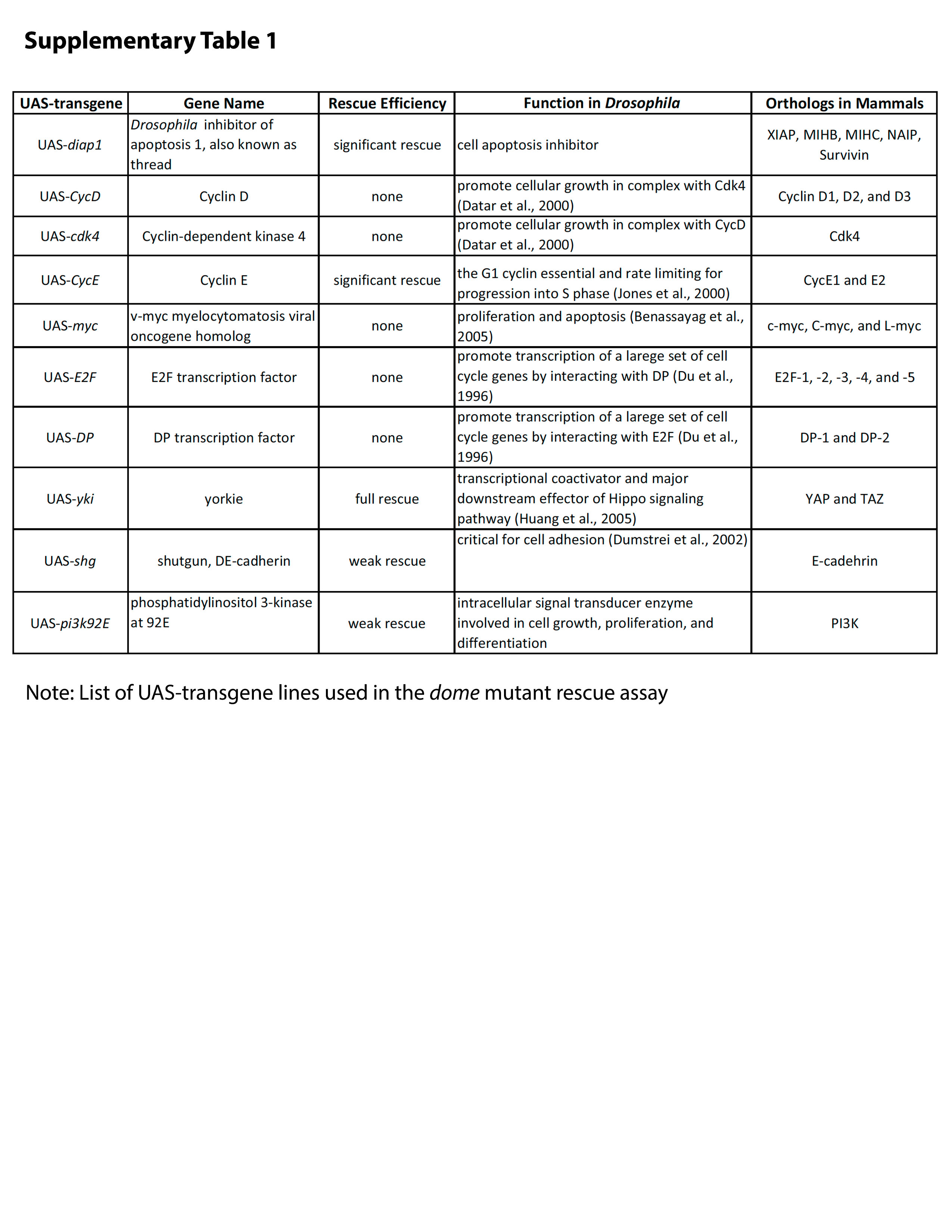
